# Thermoregulatory roles of preoptic area histamine H_1_ receptor-expressing neurons

**DOI:** 10.64898/2026.09.22.753241

**Authors:** Ramón A. Piñol, Arianna Valeri, Cuiying Xiao, Colleen K. Hadley, Abigail I. Goldschmidt, Damien Kerspern, Soumya Kulkarni, Oksana Gavrilova, Andrew Lutas, Michael J. Krashes, Marc L. Reitman

## Abstract

The preoptic area (POA) is a key regulator of body temperature, orchestrating behavioral and physiological thermoregulatory functions, including fever, torpor, and homeostatic responses to ambient temperature. Histamine regulates body temperature across animal taxa, including mammals, lower vertebrates, and invertebrates. *Hrh1*, which encodes the histamine H_1_ receptor, is expressed in several neuronal populations in the POA with limited overlap with other POA thermoregulatory neurons. Acute activation of POA^Hrh1^ neurons increased body temperature and physical activity. Acute inhibition attenuated the dark phase increase in body temperature without affecting physical activity. There was no effect of inhibition on body temperature during the light phase, fever, cold exposure, or warm exposure. Assessing behavioral thermoregulation, we found that POA^Hrh1^ neurons stimulated nest building in both sexes, possibly contributing to cold-induced nesting behavior. Through projections to the arcuate, dorsomedial hypothalamic and tuberomammillary nuclei, POA^Hrh1^ neurons can increase body temperature and physical activity. Overall, POA^Hrh1^ neurons participate in several thermoregulatory functions, including daily body temperature rhythm regulation, physical activity, and nest building behavior.

## INTRODUCTION

Tight regulation of body temperature (Tb) is a defining feature of homeothermic animals. The first line of Tb regulation is motivated behavior to modify the thermal environment, which in mice includes, nest building, huddling, and location preference. Physiological processes include regulation of vasoconstriction in superficial tissues and thermogenesis. The latter is energetically more costly than vasomotion and behavioral adaptations. Many factors contribute to selection of the defended Tb, sometimes called a set point. These include light/dark phase, sleep/wake state, nutritional status, and environmental temperature. The hypothalamus’ preoptic area (POA) is a key node in neural circuits controlling thermoregulatory behavior and physiology ^1, 2^. Many monogenic POA neuron populations have been probed for their role in thermoregulation, with most decreasing Tb on activation ^3^ and only a few increasing Tb ^4, 5^. Dedicated glutamatergic POA projections to the dorsomedial hypothalamus (DMH) are a principal neuronal pathway for heat generation via sympathetic activation of brown adipose tissue ^6^. Parallel, there are also glutamatergic POA populations with projections to the DMH that decrease Tb, playing a role in torpor and heat defense ^7^. Many other pathways and neurotransmitters, some not yet characterized, also contribute to Tb regulation.

In the central nervous system histamine is a neurotransmitter produced by histidine decarboxylase-expressing (HDC) neurons, which are located in the subdivisions of the tuberomammillary nucleus in the posterior hypothalamus ^8, 9, 10, 11^. Higher extracellular histamine levels in the brain are associated with vigilance and wakefulness ^12, 13, 14^. Early neuroanatomical studies have associated histamine in the hypothalamus with the regulation of Tb ^15^. Subsequent studies identified a conserved connection between circadian rhythm, Tb regulation, and histamine ^12, 16^. In the POA, histamine 1 receptor (H_1_R) activation can increase Tb ^17^ and increased extracellular histamine levels correlate with wakefulness ^12, 18^.

Mice are nocturnal and have a circadian rhythm for Tb and physical activity: during the dark period, mice are awake more often, eat more, are more physically active, and have a higher average Tb than during the light period ^19, 20^. The higher average dark phase Tb is the result of two independent effects. First, the Tb is ∼0.7°C higher in the dark (vs light) photoperiod. Separately, the Tb is ∼0.9°C higher during the active/awake (vs resting/sleep) state, which is more frequent during the dark period ^21^. Several hypothalamic areas have been associated with circadian Tb regulation, including the suprachiasmatic nucleus (SCN), POA, DMH, and arcuate nucleus (Arc) ^22^.

Here we investigate the role of POA histamine H_1_ receptor-expressing (POA^Hrh1^) neurons in thermoregulatory functions and energy metabolism. We demonstrate that POA^Hrh1^ neurons are one of the few POA populations that can increase Tb when activated and regulate nest building behavior.

## RESULTS

### Systemic administration of Hrh1 antagonist modestly reduces Tb

Since histamine signaling and Hrh1 are associated with the regulation of Tb, we examined the Tb responses to systemic administration of pyrilamine, a brain-penetrant H_1_R antagonist. Pyrilamine reduced Tb in female mice during the light phase (sFig 1C) and at the onset of the dark phase (sFig 1G), but not in males (sFig 1A,E). H_1_R antagonism slightly reduced physical activity in both sexes and during both phases (sFig 1B,D,F,H). Mice lacking Hrh1 (Hrh1^-/-^ mice) do not have a reduced Tb or altered physical activity (sFig 1I-L).

Having established a role for H1R in Tb regulation, we generated a Hrh1-Cre knock-in mouse line and used it to investigate how Hrh1-expressing neurons regulate Tb and energy metabolism. When bred to a GFP-reporter line (to get Hrh1-Cre;Ai6 mice), there was widespread labeling in cortical layers II/III, IV, and VI, lateral septal nucleus, claustrum, dorsal endopiriform nucleus, and multiple preoptic area regions including, MnPO, VMPO, MPA, VLPO, and LPO, replicating the reported *Hrh1* mRNA expression pattern ^23^ (sFig 1M). There was also labeling in the lateral striatum, consistent with transient Hrh1 expression at a point during development ^24^.

We focused on the Hrh1-expressing neurons in the preoptic area (POA^Hrh1^, including MnPO, VMPO, MPA, VLPO, and LPO), since preoptic area neuronal populations regulate autonomic and behavioral aspects of body temperature control. Systemic treatment with the H_1_R antagonist rapidly reduced the light phase neuronal activity of POA^Hrh1^ neurons in awake mice (sFig 1N; sFig 2A). This demonstrates that POA^Hrh1^ neurons are active during the light phase and deserving of functional study.

### Chemogenetic activation of POA^Hrh1^ neurons increases body temperature, energy expenditure, and physical activity

To determine whether activation of POA^Hrh1^ neurons regulates Tb, we expressed the Gq DREADD, hM3Dq (Fig 1A,B; sFig 2B). Like hM3Dq, H_1_R couples to Gq ^13^. Chemogenetic POA^Hrh1^ neuron activation in the light phase increased total energy expenditure (TEE), Tb, and physical activity in male mice (Fig 1C,D,F). These increases are separate from those caused by handling and dosing the mice, which last about an hour^25^. The TEE and Tb elevations were long-lived, persisting through the following dark period. The increase in physical activity was of shorter duration, closer to the ∼10 h pharmacokinetic window of CNO ^26^. The prolonged duration of the POA^Hrh1^ neuron-mediated Tb and TEE increases contrast with the shorter ∼4-6 h increases produced by activation of DMH^Brs^^3^ neurons ^25^. Chemogenetic POA^Hrh1^ stimulation in females produced similar Tb and activity responses as in males (sFig 3A,B).

**Fig 1.**
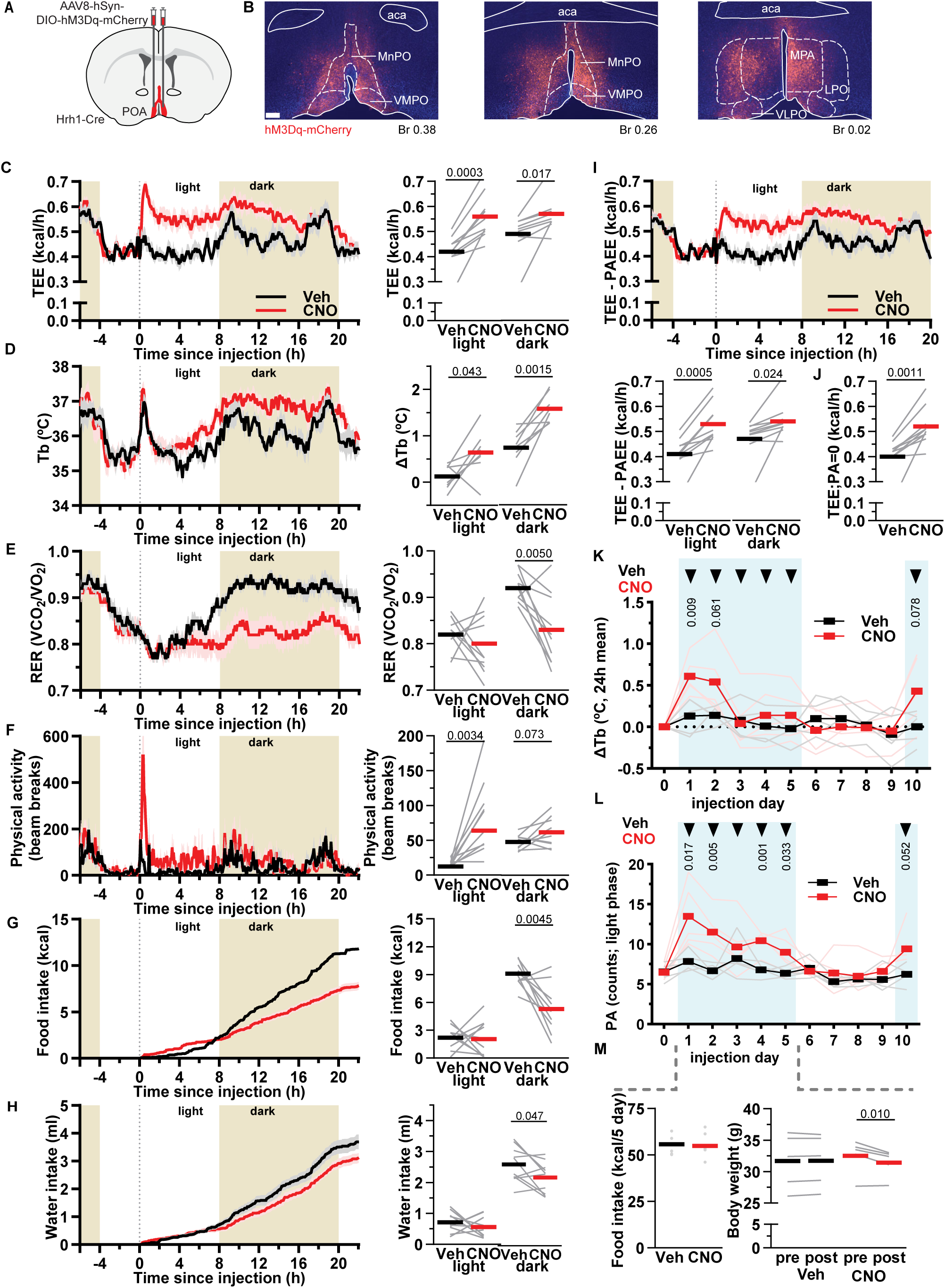
POA^Hrh1^ neuron activation increases body temperature, total energy expenditure, and physical activity. A) Schematic of virus injection and B) hM3Dq-mCherry expression in POA. Scale bar is 200 µm. C) TEE, D) Tb, E) RER, (F) physical activity, G) food intake, and H) water intake response (mean ± SEM) to vehicle (Veh) or CNO injection at t=0 (dotted line) in male mice (n = 10). Right side, quantification of mean response (bar) and individual mice (gray lines) during light (0 to 8 h) and dark (8 to 20 h). TEE, RER, and activity are the mean during the interval; ΔTb is the interval mean minus baseline (-2.5 to -0.5 h); food and water intake are cumulative intake during the interval. I) TEE response with the PAEE subtracted, otherwise as in C. J) TEE during time intervals with no physical activity between 0 and 20 h from injection. Group mean (bar) and individual mice means (gray lines). K) ΔTb (24 h mean) and L) physical activity (light period mean) response to daily repeated injections, followed by four days rest before final injection. Individual mice (n = 5 males per treatment) in lighter colors. Tb data normalized to day 0. M) Food intake and body weight over the five treatment days. P values are paired (C-J,M-right) or unpaired (K,L,M-left) t-tests. TEE: total energy expenditure; RER: respiratory exchange ratio; PA: physical activity; PAEE: physical activity energy expenditure.

While increased physical activity accompanies the increases in TEE and Tb, three lines of evidence demonstrate that the TEE and Tb increases are at least partially independent of physical activity: 1) the TEE and Tb increases last longer than the activity increase, 2) the calculated energy cost of physical activity ^27^ explains only 24% of the TEE increase (Fig 1I), and 3) the TEE was increased even when mice were not physically active (Fig 1J).

Aside from a role in Tb/TEE regulation, histaminergic neurons and H_1_R regulate food intake ^28,29^. During the first 8 hours after CNO dosing (light phase), chemogenetic activation of POA^Hrh1^ neurons did not affect food intake, but by 22 hours cumulative caloric intake was reduced by 31 ± 13% (Fig 1G). The delayed food intake suppression effect falls outside of the expected pharmacokinetic window of CNO activation of POA^Hrh1^ neuron and is possibly due to downstream circuitry. Water intake was slightly reduced (13 ± 8%) by the end of the dark period, possibly reflecting prandial drinking (Fig 1H). The respiratory exchange ratio (RER) was reduced from the onset of the dark cycle (Fig 1D), indicating increased fatty acid usage, likely a consequence of the increased TEE combined with the reduced food intake.

We next examined subchronic POA^Hrh1^ neuron stimulation with daily CNO injections for 5 days. The Tb increase attenuated after the second dose and was restored after four days off drug (Fig 1K). In contrast, physical activity remained increased during repeated CNO dosing (Fig 1L; day 1 is not significantly different from day 5 with ANOVA multiple comparisons p = 0.14). Body weight was reduced by 1.1 ± 0.3 g in the CNO group after the 5 day treatment (Fig 1M). Daily CNO treatment did not affect cumulative food intake (Fig 1M). The different attenuation characteristics of Tb and physical activity confirm that they can be independent.

These experiments show that acute activation of POA^Hrh1^ neurons robustly increases TEE, Tb, and physical activity with a prolonged duration in both sexes. Most of the TEE increase is independent of physical activity. The Tb effects attenuated over 5 days of dosing, suggesting that chronic POA^Hrh1^ neuron activation may have a modest impact on energy balance.

### POA^Hrh1^ neurons regulate dark period Tb

To examine the necessity of POA^Hrh1^ neurons in Tb regulation, we virally expressed the inhibitory chemogenetic receptor hM4Di (Fig 2A; sFig 2C). POA^Hrh1^ neuron inhibition during the light period had no effect on Tb or physical activity (Fig 2B,C; sFig 4A). In contrast, inhibition during the dark period significantly reduced Tb (by 0.24 ± 0.10 °C in males, 0.23 ± 0.12 °C in females) (Fig 2D; sFig 4B). The dark phase Tb decrease was due to a reduced resting Tb (approximated by the 5^th^ percentile Tb), while the active Tb (approximated by the 95^th^ percentile Tb) was not affected (sFig 4B). The dark phase activity, both total and resting (percent of time with no physical activity) were not affected by POA^Hrh1^ neuron inhibition (Fig 2E; sFig 4B). This suggests that POA^Hrh1^ neurons contribute to maintaining Tb during the dark phase.

**Fig 2.**
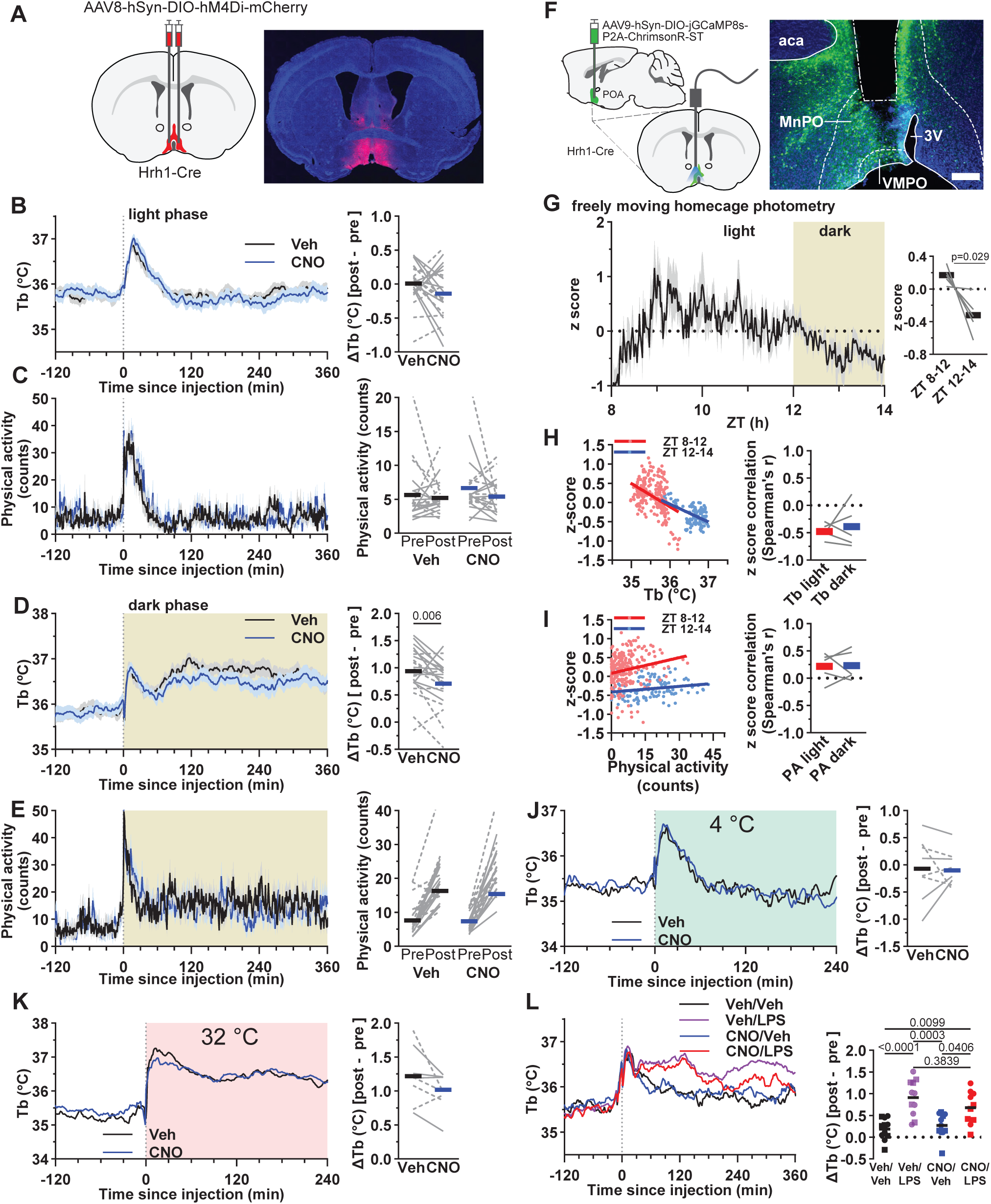
Chemogenetic inhibition of POA^Hrh1^ neurons reduces dark period Tb increase, but not light period, acute cold, or fever Tb. A) Schematic of virus injection in POA and example of hM4Di expression. B-E) Tb and physical activity response (mean ± SEM) to Veh and CNO injection at t=0 (dotted line) during the light (B,C) or at onset of dark (D,E) phase in male and female mice (n = 22). F) Schematic of virus injection strategy and optic fiber placement for measuring the Ca^2+^ response of POA^Hrh1^ neurons (left). Example of GCaMP expression and optic fiber placement (right). G) Mean z-score of Ca^2+^ fiber photometry recording (±SEM) of POA^Hrh1^ neurons in male mice (n = 5) from ZT8 to 14 (left). Quantification (right): mean (bar) and individual mice (gray lines). H, I) Left: Correlation of z-score with Tb (H) and physical activity (I). Right: Quantification: mean (bar) and individual mice (gray lines). J-L) Tb response to chemogenetic inhibition (J) at acute 4° C, (K) at acute 32° C, or (L) with LPS treatment. Quantification B-E,J-L): ΔTb, post (mean of 60 to 180 minutes) minus pre (mean of -150 to - 30 minutes); physical activity, pre is mean of -150 to -30 minutes and post is mean of 60 to 180 minutes. Mean (bar) and individual mice (males: uninterrupted lines or squares; females: dotted lines or circles). In L, t=0 aligned with second dosing. P value, in B-E, G-K is paired t-test in and, in L, one-way ANOVA with Tukey’s multiple comparison test comparing all means to each other.

Therefore, we hypothesized that POA^Hrh1^ neurons are more active during the dark phase, thereby contributing to the dark phase Tb increase. To the contrary, using Ca^2+^ fiber photometry, we found lower activity during the onset of the dark phase (ZT 12-14) compared to the light phase (ZT8-14) (Fig. 2F-I; sFig 2A,D). Fos expression in POA^Hrh1^ neurons at Zt 8 vs 14 confirmed lack of activation during the early dark phase (sFig 4C). Equally unexpectedly, we found that GCaMP activity in POA^Hrh1^ neurons is inversely correlated with Tb during light and dark phases (mean Spearman: ZT8-12, r = - 0.47 ± 0.06; ZT12-14, r = - 0.39 ± 0.18), suggesting that these neurons are more active during low Tb states. We did not observe these effects in GFP control photometry mice (sFig 4D; sFig2D). The higher activation during low Tb (Fig 2H) and dark phase reduced resting Tb during chemogenetic inhibition (sFig 4B) could suggest that POA^Hrh1^ neurons contribute to maintaining Tb during periods when Tb falls below a certain threshold.

When POA^Hrh1^ neurons were inhibited and the mice were exposed to 4 °C during the light phase, we observed no effect on Tb or physical activity (Fig 2J; sFig 4F). This suggests, consistent with the results at 22 °C, that POA^Hrh1^ neurons do not appear to be required for acute cold defense thermogenesis. Similarly, POA^Hrh1^ neuron inhibition during exposure to 32 °C did not change Tb (Veh, 36.42 ± 0.21 °C; CNO, 36.48 ± 0.18 °C) or physical activity (Fig 2K; sFig 4G).

Activation of POA neurons can produce a fever response ^5^. Thus, we tested the effect of POA^Hrh1^ neuron inhibition during two experimental fever models. Inhibition of POA^Hrh1^ neurons did not affect LPS- or polyI:C-induced fever in males or females (Fig 2L; sFig 4E, H, I). Stress produces an increase in Tb and physical activity. Inhibition of POA^Hrh1^ neurons did not reduce the Tb or physical activity responses to handling and injection in either males or females (Fig 2B,C; sFig 4J).

Together, these data indicate that POA^Hrh1^ neurons contribute to the dark period Tb increase, but do not appear to be required for Tb regulation during the light period, in fever, or with handling stress.

#### Brief optogenetic activation of POA^Hrh1^ neurons drives a prolonged body temperature increase, but does not surpass a ceiling Tb

Optogenetics allows neuronal stimulation with precise temporal control without handling or disturbing the mice. We virally expressed ChR2 in POA^Hrh1^ neurons and fiber placement centered on the anterior POA (sFig 2D,E). Activation of anterior POA^Hrh1^ neurons during the light phase increased Tb and physical activity (Fig 3A; sFig 5A,B). Physical activity reached maximum levels immediately, while the Tb increase was slower due to the heat capacity of the mouse (sFig 5A,B). The greatest Tb and physical activity responses were achieved using a stimulation frequency of 10-20 Hz (sFig 5C,D). For comparison with POA^Brs3^ neuron activation, we used the same stimulation paradigm (5 cycles of 20 minutes followed by 60 minutes off) ^4^. Physical activity rapidly returned to baseline while Tb remained above baseline after 60 minutes, causing upward drift of each cycle’s pre-stimulation ‘baseline’ Tb (quantified as the ‘Off’ interval; Fig 3B,C). This contrasts with POA^Brs3^ neurons, which completely returned to baseline by 60 minutes after each cycle. Using a single 40-minute POA^Hrh1^ stimulation, some Tb increase persisted for hours, although determining a precise duration would require a much larger cohort size (sFig 5E). Consistent with the chemogenetics results, the optogenetically induced physical activity increase matched the stimulation duration while the Tb remained elevated long after stimulation termination (Fig 3C; sFig 5F). Stimulation duration did not affect the kinetics of the return of physical activity to resting, while 20 and 40 min stimulation may delay the return of Tb to baseline (sFig 5G,H).

**Fig 3.**
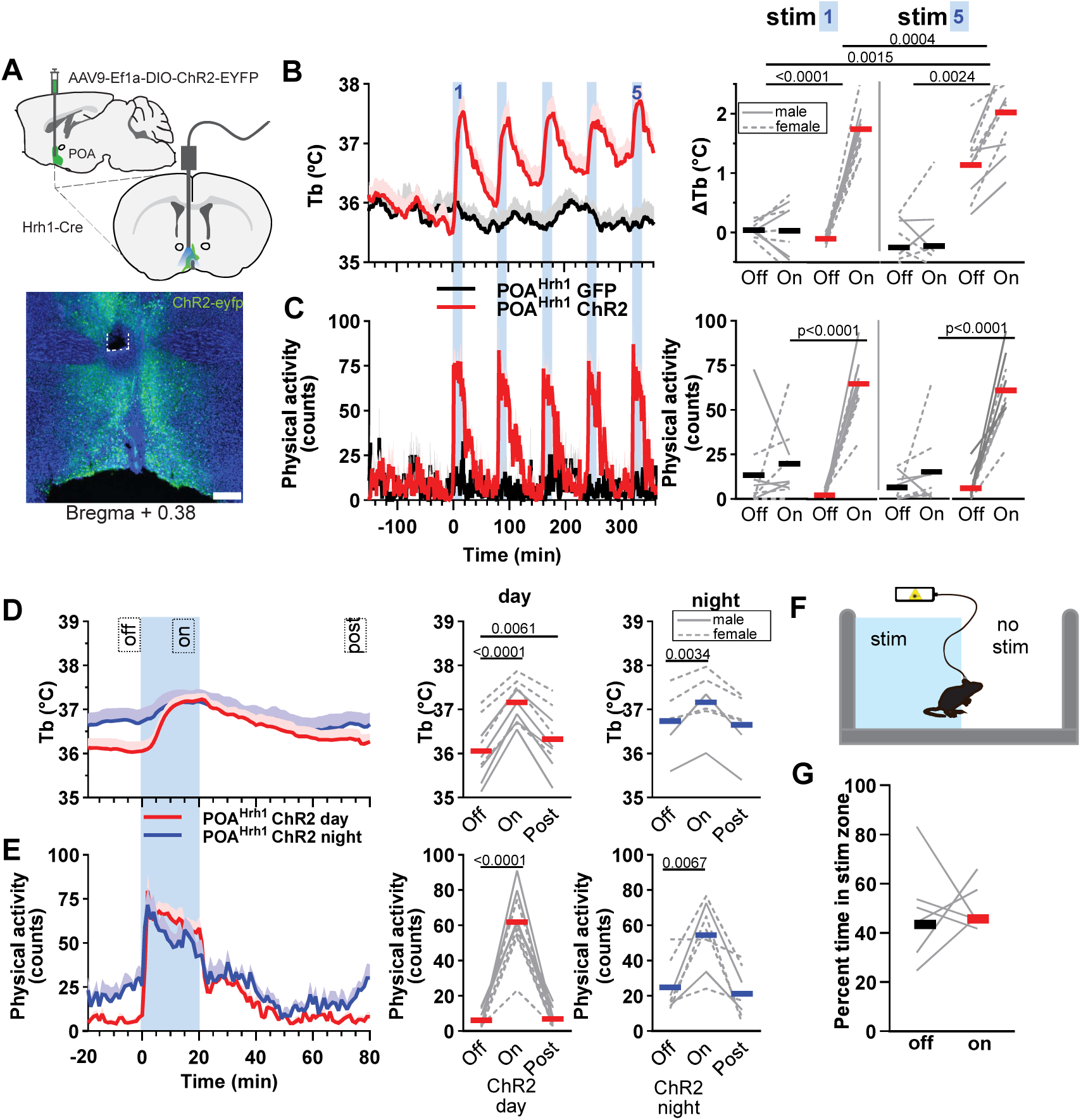
Optogenetic POA^Hrh1^ neuron activation increases body temperature and physical activity to similar absolute levels during light and dark period. A) Schematic of virus injection and optic fiber placement and (bottom) example showing ChR2-EYFP expression and fiber site (dotted line). Scale bar is 200 µm. B) Tb and (C) physical activity response (mean + SEM) to five cycles of 20 minutes laser on (blue bar) followed by 60 minutes laser off during light phase (left side) in male (n = 5) and female (n = 5) mice. Right side: quantification of the first and fifth cycles with group mean (bar) and individual mice (gray lines). ΔTb: ‘Off’ is the mean of the 10 minutes immediately preceding stimulation and ‘On’ is mean of laser minutes 10-20 minus baseline (-20 to 0 min). Physical activity: ‘Off’ is the mean of the 10 minutes immediately preceding stimulation and ‘On’ is the mean of laser minutes 0-20. D) Mean Tb and (E) physical activity responses of the five stimulation cycles (like in B) during light and dark phases in male (n = 2 to 5) and female (n = 4 to 5) mice. Right side, quantification: ‘Off’ is the mean of minutes -10 to 0; ‘On’ is the mean of minutes 10-20 for Tb and 0-20 for physical activity; ‘Post’ is the mean of minutes 70-80. Means (bar) and individual mice (gray lines) are presented. F) Schematic of real-time place preference experiment. G) Percent time spent on the stimulation side (bar: mean ± SEM; dots: individual mice) in control experiment with laser off (‘off’) and in experiment with laser on (‘on’). B-C: p values are from one-way ANOVA with Tukey’s multiple comparison test comparing all means to each other; ΔTb, all Off/On pairs (Off minus On) are compared; physical activity, all ‘Off’ and ‘On’ conditions are compared to each other. Only relevant significant values shown for clarity. D, E: p values are from one-way ANOVA with Dunnett’s multiple comparisons test comparing to ‘Off’; Tb statistics done on ΔTb where the baseline (mean of minutes -20 to 0) is subtracted from ‘Off’ (the mean of minutes -10 to 0), ‘On’ (minutes 10-20) and ‘Post’ (minutes 70-80); for physical activity ‘Off’ is the mean of minutes -10 to 0, ‘On’ of minutes 0-20 and ‘Post’ of minutes 70-80.

In the dark phase, baseline Tb and physical activity levels are higher than in the light phase. Optogenetic stimulation during the dark phase of both males and females produced a peak Tb that matched the light phase peak, which is a smaller increase from the higher dark phase baseline (Fig 3D,E). This indicates that activation of POA^Hrh1^ neurons does not cause Tb to surpass a ceiling – typically when thermogenesis is experimentally stimulated, Tb does not increase more than 1-2 °C, because heat dissipation mechanisms let go of produced warmth. Therefore, our results suggest that the activation of POA^Hrh1^ neurons does not restrict heat dissipation, like for instance during high fevers. Tb did not remain elevated after stimulation termination like it did during the light phase optogentic stimulation.

To examine the valence of POA^Hrh1^ neuron stimulation, a real-time place preference experiment was performed. No side preference was observed with optogenetic stimulation (Fig 3F,G), suggesting that POA^Hrh1^ neuron activation has neither positive nor negative valence in this assay.

There was no difference in food intake during or after optogenetic stimulation (sFig 5I,J), similar to what was observed in the early time points with chemogenetic activation.

Thus, optogenetic activation of POA^Hrh1^ neurons produces a Tb increase that far outlasts the duration of stimulation, suggesting the involvement of downstream circuitry. The Tb reached similar levels during the light and dark periods, suggesting that POA^Hrh1^ neuron stimulation does not surpass a ceiling Tb.

### Stimulation of the Vgat-expressing subset of POA^Hrh1^ neurons increases Tb

The POA is a heterogeneous region. In a POA scRNAseq dataset ^30^, 64% (848/1318) of the neurons that expressed *Hrh1* fell in inhibitory (*Vgat*-expressing) and 36% (470/1318) in excitatory (*Vglut2*-expressing) clusters, but 9% of neurons expressed both *vgat* and *vglut2* (Fig 4A; sFig 6). The POA^Hrh1^ clusters showed limited overlap with populations whose role in thermoregulation has been examined ^3, 4, 7, 31, 32, 33, 34, 35, 36, 37, 38^ (Fig 4A). Since both POA^Hrh1^ and POA^Brs3^ neurons increase Tb when activated, the extent of *Hrh1*/*Brs3* overlap was examined. Only 15% of Hrh1-Cre neurons also expressed a Brs3-FlpO-dependent reporter (Fig 4B,C). This is consistent with the 4-10% of POA *Hrh1*-expressing neurons co-expressing *Brs3* in ^30, 39^ (Fig 4D).

**Fig 4.**
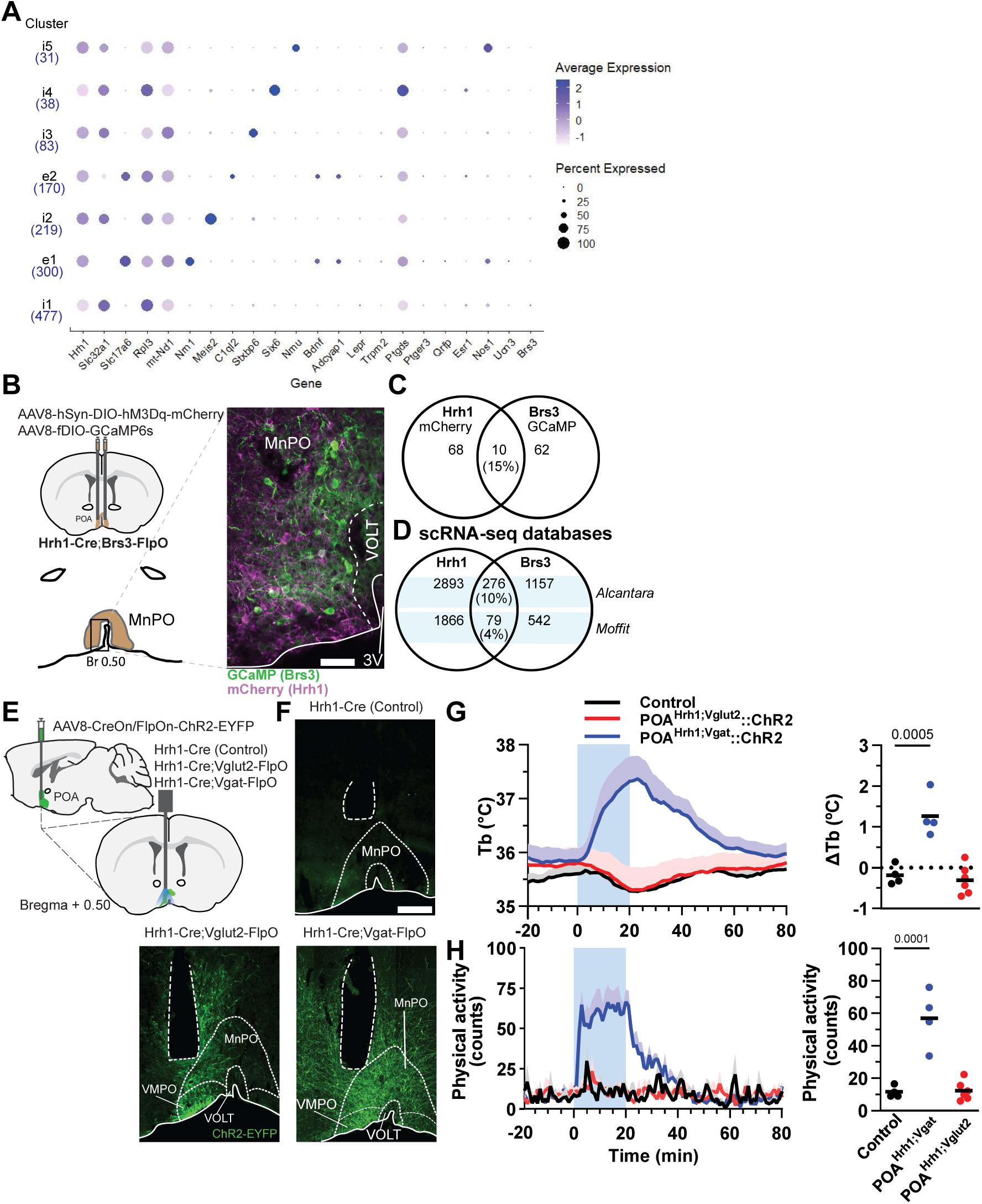
POA^Hrh1^ neurons form multiple clusters and POA^Hrh1;Vgat^ neurons mediate Tb phenotype. A) Multiple clusters of the Hrh1-expressing POA neurons; analysis of ^30^ data. Clusters expressing Slc32a1 are marked as inhibitory (‘i’) and those expressing Slc17a6 excitatory (‘e’). Number of cells per cluster is below cluster name. B-D) Quantification of the *Hrh1* and *Brs3* overlap in expression. B) Schematic and example of viral injection into MnPO of Cre-dependent hM3Dq-mCherry and FlpO-dependent GCaMP of Hrh1-Cre;Brs3-FlpO mice. Scale bar is 50 µm C) Quantification of MnPO neurons expressing both Hrh1-Cre and Brs3-FlpO reporters (n = 3 mice; neurons in 1 slice per mouse at Br 0.50 in MnPO counted). D) Overlap of *Brs3* and *Hrh1* expression in published POA scRNA-seq datasets ^30, 39^. E-H) Selective activation of POA^Hrh1;Vgat^ or POA^Hrh1;Vglut2^ neurons. E) Schematic of virus injection for intersectional strategy. F) Examples of ChR2-EYFP expression and optic fiber placement in three genotypes. Scale bar is 200 µm. G,H) Mean Tb and physical activity response (left side; mean + SEM) to five sequential (as in Fig 2B) 20-minute laser stimulation periods (blue bar) in male mice (n = 4 – 6 males/group). Quantification (right side): ΔTb, mean of laser minutes 10-20 minus baseline (-20 to 0 min); physical activity, mean of laser minutes 0-20; means (bar) and individual mice (dots) are presented G,H: p values are from one-way ANOVA with Dunnett’s multiple comparisons test comparing to ‘Control’.

We next used an intersectional approach to distinguish between inhibitory and excitatory POA^Hrh1^ neuron functions (Fig 4E; sFig 2F-H). Optogenetic activation of POA^Hrh1;Vgat^, but not POA^Hrh1;Vglut2^, neurons increased Tb and physical activity, (Fig 4F-H). These experiments demonstrate that inhibitory, but not excitatory, POA^Hrh1^ neurons can mediate an acute Tb and physical activity increase. This is the only reported POA Vgat-expressing sub-population that can increase Tb when activated.

### POA^Hrh1^ neurons have multiple inputs and broad projections

To understand POA^Hrh1^ neuron connectivity, we mapped projections using viral anterograde tracing (sFig 7A,B; sFig2J). There were abundant POA^Hrh1^ projections to the PVH, DMH, Arc, PAG, RPa, and VTM, in addition to local POA connections (sFig 7C,D).

To identify inputs to POA^Hrh1^ neurons, we performed monosynaptic retrograde rabies tracing (Fig 5A). Starter neurons were predominantly located in the MnPO or MPA (Fig 5B,C; sFig 7E). Local input from within the POA accounted for 36±2% of the input neurons, with the MnPO and MPA having the largest fraction. Multiple hypothalamic regions, including the Arc, VMH, LH, DMH, and PVH, provided another 32±5% of the input neurons (Fig 5D,E,G). Input from the area with most histaminergic neurons, the VTM, was modest at 0.5±0.04% (Fig 5E,G). Input regions from outside the hypothalamus included the lateral septum, BNST, and LPB (Fig 5F,G).

**Fig 5.**
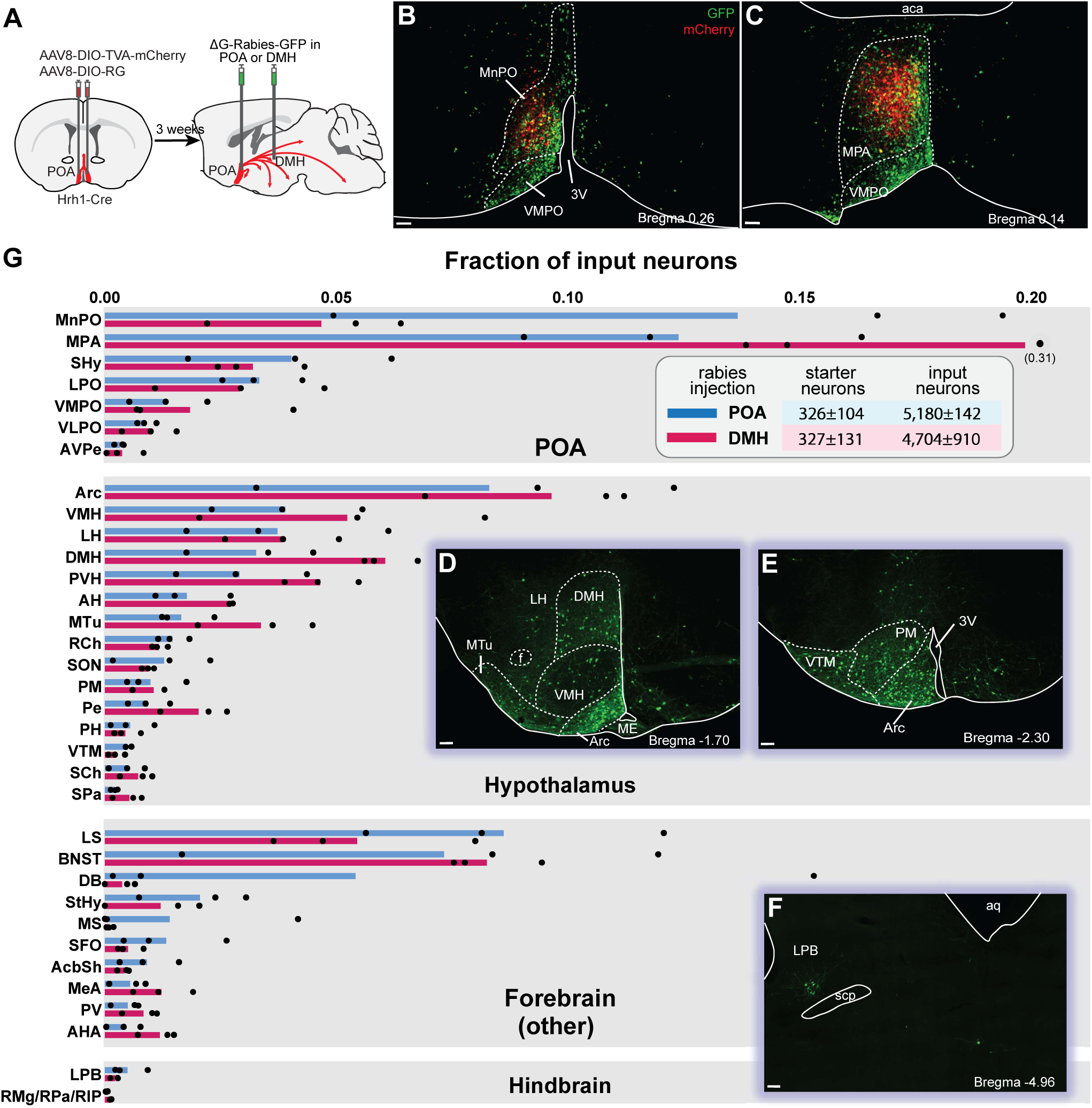
Monosynaptic retrograde tracing reveals similar input patterns to POA^Hrh1^ and POA^Hrh1^→DMH neurons. A) Schematic of virus injection strategy. Helper viruses expressing TVA-mCherry and RG were injected in POA. Three weeks later, ΔG-Rabies-GFP was injected in POA or DMH. B,C) Starter (GFP + mCherry) and POA local input (GFP only) neurons to POA^Hrh1^ neurons. D-F) input (GFP only) neurons to POA^Hrh1^ neurons. G) Quantification of input neurons to POA ^Hrh1^ (blue) and POA^Hrh1^→DMH (red) neurons as fraction of total input neurons. Only areas with input neurons in all 3 mice are shown. B-F: Scale bar is 100 µm.

The DMH is a major contributor to the control of energy expenditure ^40^. Monosynaptic retrograde tracing of the DMH-projecting POA^Hrh1^ neurons was compared to tracing from POA^Hrh1^ neurons nonselectively. This revealed a similar number of starter and input neurons (Fig 5A; sFig 7E). Additionally, the patterns of inputs to DMH-projecting vs total POA^Hrh1^ neurons were remarkably similar (Fig 5G). Thus, DMH-projecting POA^Hrh1^ neurons do not appear to be a selective subset of the general POA^Hrh1^ neuron population.

We next investigated which of the POA^Hrh1^ output pathways drive increased Tb and physical activity. After expressing ChR2 virally in POA^Hrh1^ neurons, optogenetic stimulation of POA^Hrh1^ axons in the VTM, Arc, or DMH each increased Tb and physical activity similarly to direct POA^Hrh1^ cell body activation (Fig 6A-D; sFig 2K,L). We determined that the POA^Hrh1^→DMH neurons collateralize widely, including to the Arc, VTM, PVH, RPa, and PAG and other areas (sFig 7A,B). One interpretation is that target neurons in VTM, Arc, and DMH can each increase Tb and physical activity. Alternatively, the increase in Tb and physical activity could be due to optogenetic terminal stimulation antidromically activating other collateral targets of the POA^Hrh1^ neurons ^41^. Of note, both POA^Hrh1;Vgat^ and POA^Hrh1;Vglut2^ neurons had axonal projections to DMH, Arc and VTM (sFig 8C).

**Fig 6.**
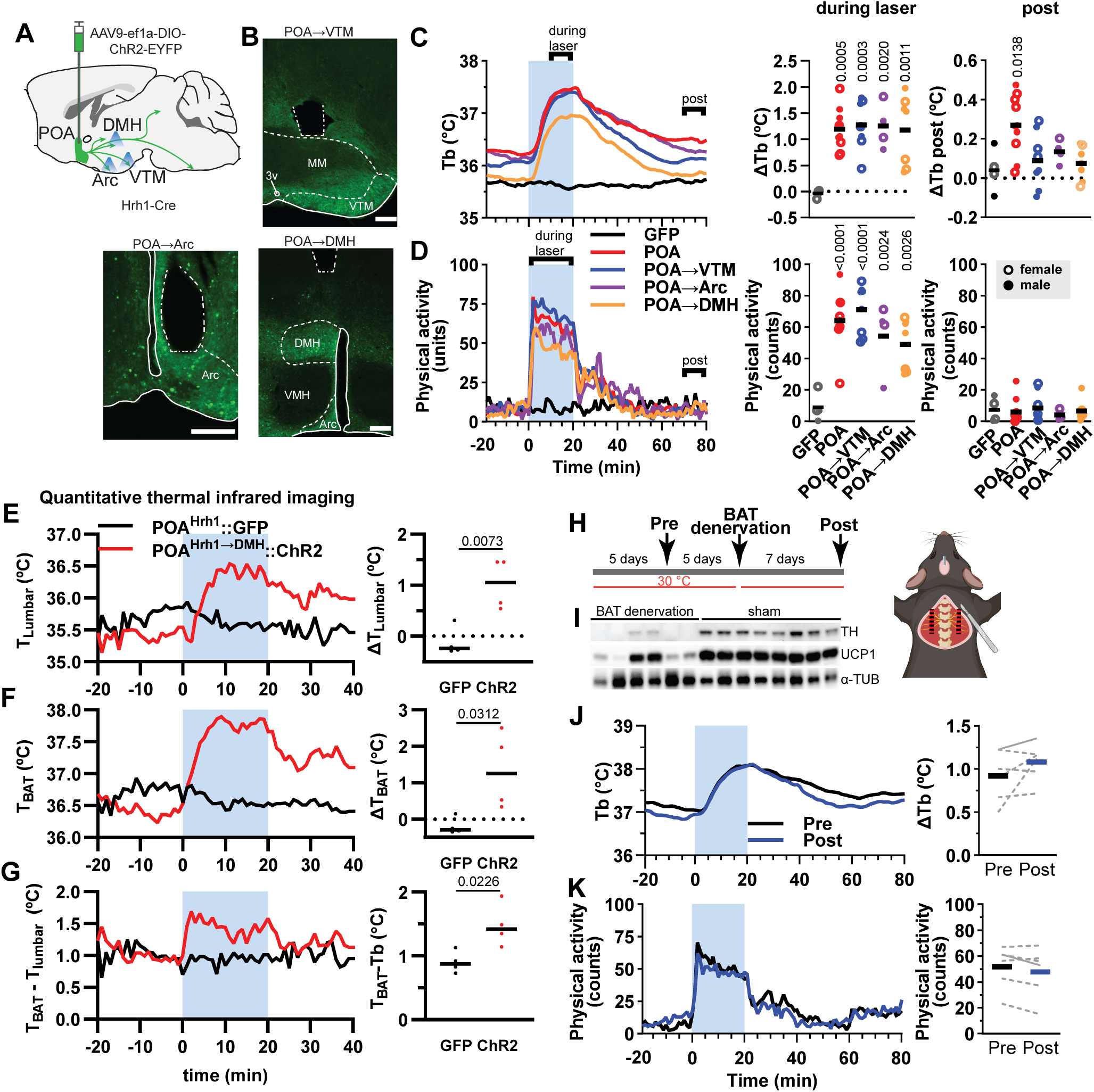
Optogenetic stimulation of axons in multiple projection areas increases Tb, which is partially mediated by BAT. A) Schematic of virus injection. Optic fibers were implanted over right DMH, Arc, or VTM. B) Examples of ChR2-EYFP expression and optic fiber placement sites in three projection areas. Scale bars are 100 µm. C,D) Mean Tb and physical activity response (left side; mean with errors omitted for clarity) to five 20-minute laser stimulation periods (blue bar) in male and female mice (n = 5-8/group). Quantification (center): ΔTb, mean of laser minutes 10-20 minus baseline (-20 to 0 min); physical activity, mean of laser minutes 0-20. Quantification (right side): ΔTb, mean of laser minutes 70-80 minus baseline (-20 to 0 min); physical activity, mean of laser minutes 70-80. Bars are means; dots, individual animals. E-G) Infrared imaging of interscapular (T_BAT_) and lumbar (T_lumbar_) skin temperature. Effect of optogenetic stimulation (blue interval) of POA^Hrh1^→DMH neurons (red, n=4) and GFP controls (black, n = 3) on T_BAT_, T_lumbar_, and the difference (T_BAT_ - T_lumbar_). Data are average of 2 trials/mouse. T_BAT_ and T_lumbar_ quantitation (right): ΔT, mean of laser minutes 10-20 minus baseline (-10 to 0 min). Experiments were performed at 25 °C. Bars are means; dots, individual animals. H) BAT denervation surgery schematic and optogenetic stimulation timeline of POA^Hrh1^ ChR2 mice. Animals were maintained at 30 °C (including during the experiment) except for during the surgery. I) Western blot for tyrosine hydroxylase (TH), uncoupling protein-1 (UCP1) and α-tubulin (α-TUB). J,K) Tb and physical activity responses (mean with errors omitted for clarity) to optogenetic stimulation ‘Pre’ and ‘Post’ denervation, analyzed as in Fig 5C,D (center). Bars are mean and lines individual animals (solid line, male; dashed line, female). C,D: p values are from one-way ANOVA with Dunnett’s multiple comparisons test comparing to ‘GFP.’ E-F: p values from unpaired t test. J,K: paired t test, not significant.

Interestingly, POA^Hrh1^ neurons receive input from and project to the VTM (Fig 5E,G; sFig 8C,D), which is where a cluster of HDC-expressing and thus histamine-producing neurons is located ^13^. POA^Hrh1^ neuron activation increased Fos expression in VTM^HDC^ neurons (sFig 8D-G). This suggests the possibility of a VTM^HDC^→POA^Hrh1^→VTM^HDC^ circuit.

The POA^Hrh1^ neuron mapping studies identify wide inputs and outputs, including a potential feedback/feedforward circuit with the VTM. This reciprocal connectivity can speculatively contribute to the role of histamine in arousal or vigilance ^14, 42^. POA^Hrh1^ neurons projecting to the DMH can increase Tb and physical activity and have widespread collaterals, which may contribute to these effects.

### POA^Hrh1^ neurons increase Tb through brown adipose tissue activation

In mice, activation of brown adipose tissue (BAT) is a major heat source for maintaining Tb, including during cold, fever, social stress, and the active/awake state ^21, 43, 44^. We studied the effect of POA^Hrh1^→DMH neuron activation, finding that optogenetic stimulation increased both Tb and T_BAT_ (Fig 6E-G). The interscapular BAT warmed up ∼0.5 °C more than the body core, indicating that POA^Hrh1^→DMH neurons increased BAT heat generation, which contributes to the Tb increase.^45^

We next asked if BAT is necessary for POA^Hrh1^ neuron-stimulated heat generation. The five sympathetic nerve bundles to interscapular BAT were cut bilaterally (Fig 6H). Denervation was confirmed by a strong reduction of BAT tyrosine hydroxylase protein, a marker for sympathetic innervation, as well as reduced UCP1 protein and an increase in BAT adipocyte fat content (Fig 6I; sFig 8H). We compared the Tb and physical activity response to optogenetic activation of POA^Hrh1^ neurons before and after the denervation and found no change (Fig 6J,K). These results suggest that POA^Hrh1^ neurons can increase Tb redundantly, via interscapular BAT and additional mechanisms (e,g, shivering, muscle thermogenesis, vasomotion).

### Prolonged Tb increase beyond activation of POA^Hrh1^ neurons is not due to persistent cell body activation

Optogenetic activation has the potential to increase circuit level activation ^46^. Therefore, a possible explanation for the Tb increase persisting long after chemogenetic or optogenetic stimulation is prolonged activation of the POA^Hrh1^ neurons, although it has not been reported that ChR2 stimulation produces persistent firing after light offset. To examine this, we virally expressed soma-targeted ChrimsonR for optogenetic activation and GCaMP8s to monitor intracellular Ca^2+^ levels (Fig 7AB; sFig2A,D,M). Optogenetic stimulation of the POA^Hrh1^ neurons increased Ca^2+^ levels time-locked with the laser pulses and, immediately following, reduced Ca^2+^ levels transiently below baseline, suggesting that the neurons were active prior to optogenetic stimulation. However, there was no increase in Ca^2+^ levels at later times (Fig 7C,D). As a comparator, POA^Brs3^ neurons also showed both the post-stimulation undershoot and no late Ca^2+^ changes.

**Fig 7.**
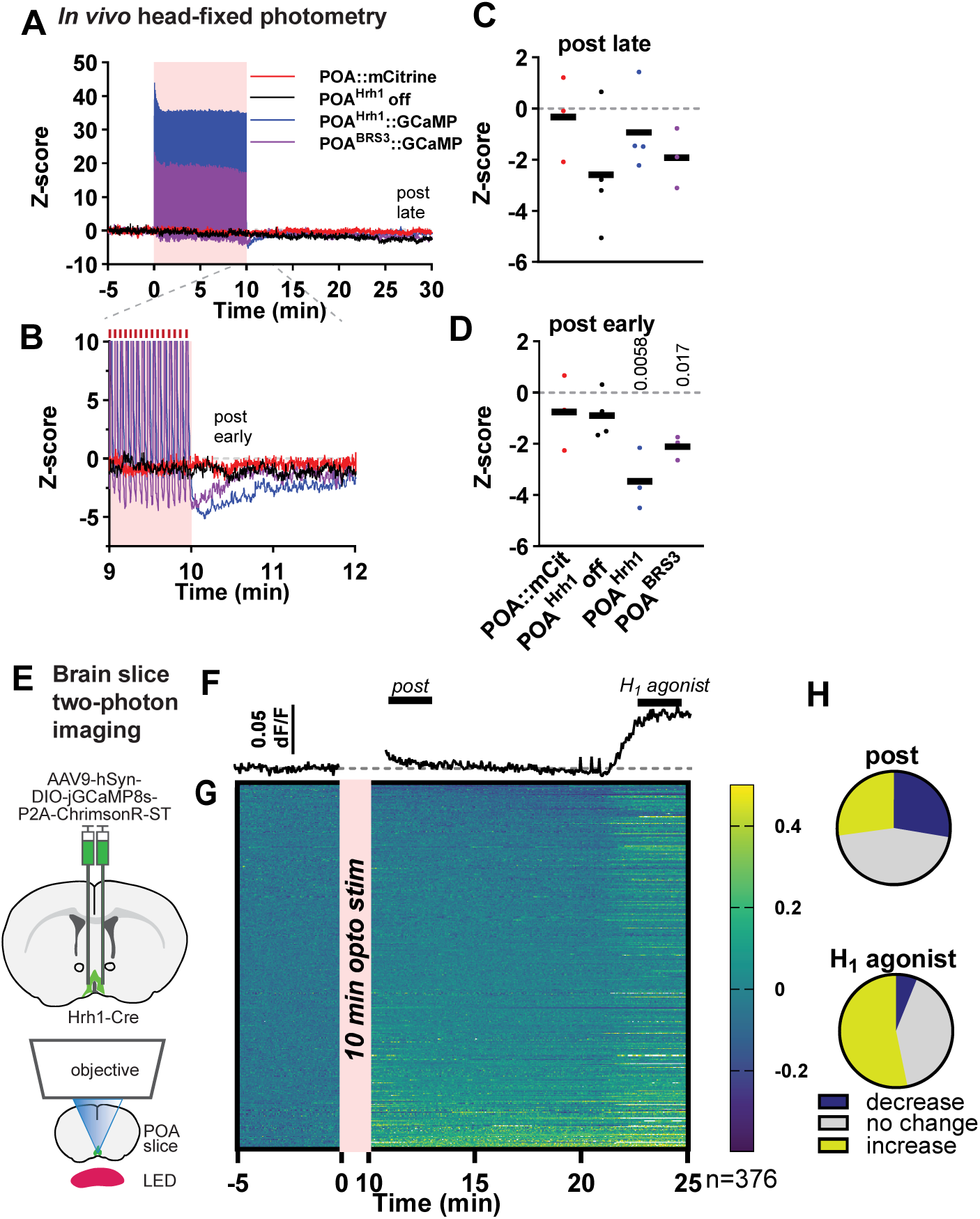
No prolonged neuronal activity in POA^Hrh1^ neurons after stimulation. A) Mean of Ca^2+^ fiber photometry recording response (SEM omitted for clarity) of POA^Hrh1^::mCitrine (red), POA^Hrh1^ with laser off (black), POA^Hrh1^::GCaMP (blue), and POA^Brs3^::GCaMP (purple) male mice to 10 minutes optogenetic stimulation (light red interval). N = 3-4/group. B) expanded view of graph in panel A around end of optogenetic stimulation. C, D) Quantification: bars are means and dots individual mice (n = 4 male mice/group); ‘Post early,’ mean of minute 10-11; ‘Post late,’ mean of minutes 25-30. E) Schematic of virus injection strategy and 2-photon POA slice imaging. F,G) mean and individual cell Ca^2+^ response to 10 minutes of optogenetic stimulation (light red interval) followed by H_1_R agonist PEA (100 µM). Heatmap sorted by response from 10-19 minutes. H) Fraction of POA^Hrh1^ neurons with a Ca^2+^ response after the end (mean of minutes 11-13) of optogenetic stimulation (top) and (bottom) to PEA (the mean of minutes 23-25 minus pre-agonist baseline (the mean of minutes 17-19)). C,D): p values are from one-way ANOVA with Tukey’s multiple comparison test comparing all means to each other.

We also imaged POA^Hrh1^ neurons in brain slices from mice injected with the ChrimsonR-GCaMP8s virus (Fig 7E), allowing us to observe individual neuron responses rather than a bulk Ca^2+^ signal. After optogenetic stimulation, 27% of the neurons increased Ca^2+^ levels, while 28% were below the pre-stimulation baseline (Fig 7F-H). Taken together, these experiments suggest that the prolonged Tb increase is not due to continued POA^Hrh1^ neuron activation and presumably involves synaptic plasticity of downstream neurons. However, we cannot exclude that a subset of persistently active neurons contributes to the prolonged Tb increase.

We also tested the Ca^2+^ response of Hrh1-Cre neurons to pharmacological H_1_R activation. When the POA slices were treated with 2-Pyridylethylamine (PEA), a selective H_1_R agonist, Ca^2+^ levels increased in 53% of the neurons (Fig 7H-J). These agonist and antagonist responses (sFig 1N) demonstrate that the H_1_R receptors are functional in the POA^Hrh1^ neurons.

### POA^Hrh1^ neurons bidirectionally regulate nest building, but not thermal preference

POA neurons are activated by acute exposure to cold ambient temperatures ^47^. Exposure to 4 °C induced Fos expression in 32 ±1.6 % of MnPO^Hrh1^ neurons, compared to 6 ±0.9 % at 30 °C (Fig 8A-C; sFig 9A). 30 °C during the light phase is close to the thermoneutral point of mice and they do not use any thermogenesis to regulate Tb ^48^. There was greater Fos/Hrh1 colocalization in the MnPO than in the more caudally located MPA.

**Fig 8.**
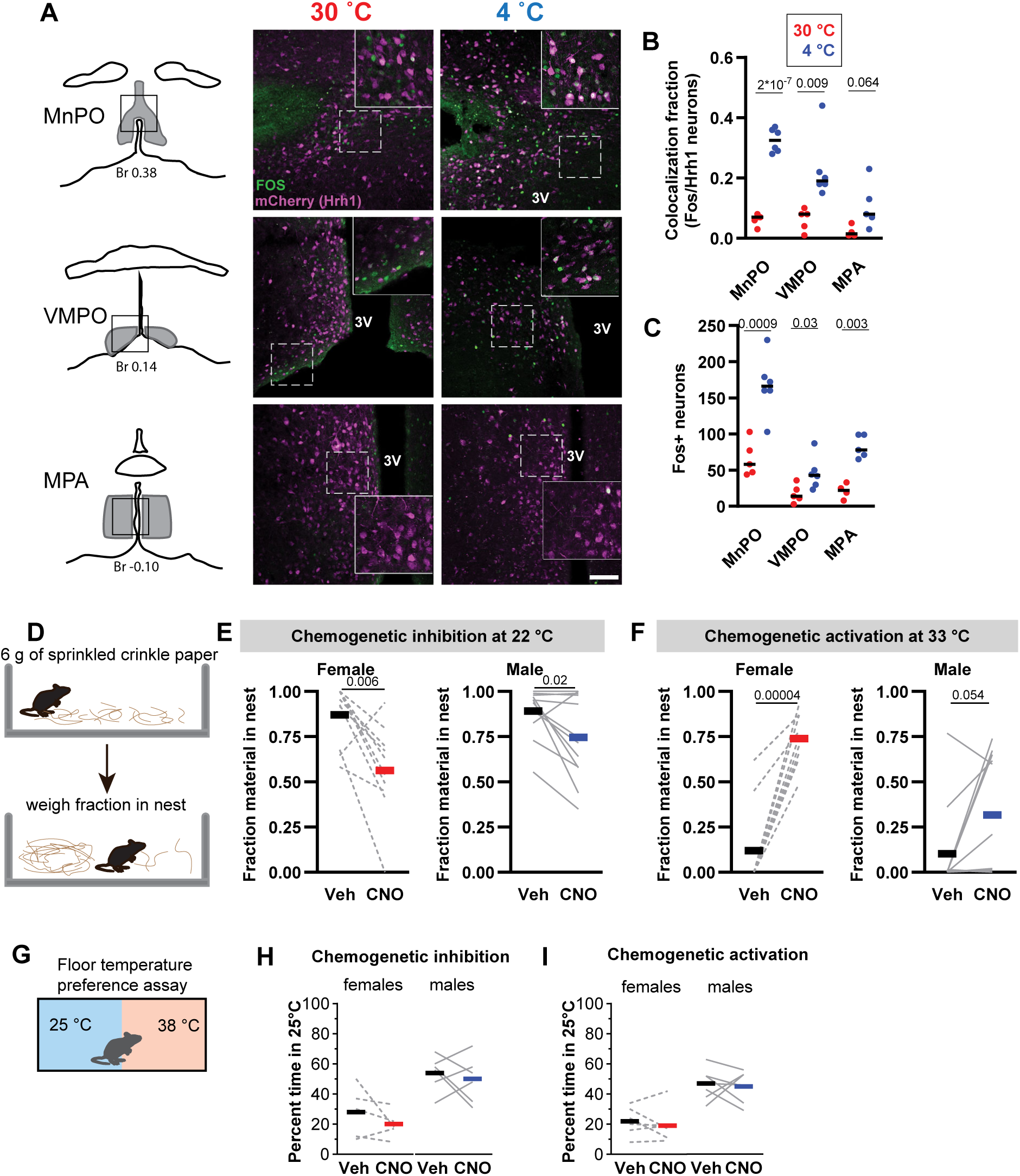
POA ^Hrh1^ neurons are cold-activated and contribute to nest-building behavior. A) Examples of Fos expression after 3 h of acute 30° C or 4° C exposure (during the light phase) in MnPO, VMPO and MPA in Hrh1-expressing neurons. Scale bar is 100 µm; inset is 2x magnification. B) Fraction of Hrh1 neurons that expresses Fos in MnPO (140 ± 14 mCherry neurons), VMPO (76 ± 8 mCherry neurons) and MPA (173 ± 25 mCherry neurons), and C) number of Fos neurons per nucleus in male and female mice (n = 6). D) Schematic of nest-building behavior assessment. E,F) Nest-building behavior in response to CNO or veh in female (n = 9-10) and male (n = 10-11) mice G) Schematic of two-temperature choice test. H,I) Mean percentage of time in 25°C in response to CNO or veh. P value is of unpaired (B,C) or paired (E,F) t-test

Next, we tested the POA^Hrh1^ neuron contributions to two thermoregulatory behaviors, nest building and thermal preference. Inhibition of POA^Hrh1^ neurons strongly reduced nest building at 22 °C (Fig 8D,E; sFig 9B-D). Conversely, mice do not usually build nests at hot ambient temperatures, however, chemogenetic activation of POA^Hrh1^ neurons caused nest building at 33 °C (Fig 8F).

Organisms seek preferred temperatures as a thermoregulatory behavior and histamine is crucial for *Drosophila* temperature preferences ^16^. However, we did not detect a change in thermal preference after either chemogenetic inhibition or activation of POA^Hrh1^ neurons in males or females (Fig 8G-I).

These experiments show that POA^Hrh1^ neurons are cold sensitive and bidirectionally regulate nest building, with activation stimulating nest building.

## DISCUSSION

We demonstrate that activation of POA^Hrh1^ neurons can increase Tb independent of physical activity. Specifically, POA^Hrh1;Vgat^ but not POA^Hrh1;Vglut2^ neurons regulate Tb; further identification of subclusters remains to be determined. While most tested monogenic POA populations lower Tb when activated, several increase Tb, including POA^Brs3^ and VMPO LPS-activated (VMPO^LPS^) populations ^4, 5^. Parenthetically, LPS inhibits EP3R-expressing neurons, which increases Tb ^31^. *Hrh1* has very limited expression in the POA^Brs3^ and VMPO^LPS^ populations, which also have different functions. Differences between POA^Hrh1^ and POA^Brs3^ neurons include that excitatory POA^Brs3^ neurons raise Tb and contribute to cold defense, while POA^Hrh1^ neurons that raise Tb are inhibitory and do not decrease Tb when inhibited during the light phase. Furthermore, chemo- or optogenetic activation of POA^Hrh1^ neurons, but not POA^Brs3^ neurons, leads to continued elevated Tb after cessation of activation. Also, VMPO^LPS^ neurons, consisting of non-overlapping VMPO^Calcr^ and VMPO^Gal^ populations, regulate fever, but POA^Hrh1^ neurons do not. While both VMPO^LPS^ populations increase Tb, VMPO^Gal^ neurons also mediate temperature preference and VMPO^Calcr^ neurons mediate immediate food intake suppression, behaviors that were not affected by POA^Hrh1^ neuron manipulations. Further work is necessary to pinpoint the neuroanatomical location within the POA nuclei of POA^Hrh1;Vgat^ neurons that increase Tb and to identify properties that define Tb-regulating POA^Hrh1;Vgat^ subclusters.

POA^Hrh1^ neurons receive input from a wide variety of brain areas, including a small fraction from the VTM. Some POA^Hrh1^ neurons have also axonal projections back to the VTM. This anatomy suggests the possibility of a positive feedback loop, with histamine from VTM^HDC^ neurons activating POA^Hrh1^ neurons through H_1_R G_q_ signaling ^12^ and the POA^Hrh1^ neurons then activating VTM^HDC^ neurons.

Stimulation of POA^Hrh1^ terminals in the VTM, DMH, and Arc each increased physical activity and Tb. Our experiments do not differentiate between the possibility of each target area driving Tb increase and antidromic collateral activation reaching the physiological output regions that drive the response ^41^. Collectively, our observations suggest that POA^Hrh1;Vgat^→DMH neurons can increase Tb. A possible pathway includes direct synaptic connection of POA^Hrh1;Vgat^→DMH neurons to local inhibitory interneurons in the DMH region ^47^. When inhibited, these inhibitory interneurons remove a brake on thermogenic DMH→RPa neurons to drive BAT thermogenesis ^2^.

While VTM^HDC^ neuron firing correlates with being awake ^49^, a causal role for these neurons in sleep/wake behavior has not been established ^50, 51, 52^. Chemogenetic activation or increasing the excitability of VTM^HDC^ neurons, did not change sleep/wake behavior, except for an increase in wakefulness after a cage change ^51, 52^. Additionally, *Hrh1^-/-^* mice have an unchanged sleep/wake pattern, but in the light period fall asleep faster after transfer to a new cage. These responses to the novelty of a new cage are likely mediated by H_1_receptor ^50^ and may reflect a state of increased vigilance ^14^.

Could there be a role for POA^Hrh1^ neurons in thermoregulation related to sleep? Mouse sleep preparation and initiation includes building and curling up in nests. In humans, gentle skin warming facilitates sleep onset and reinforces sleep maintenance (reviewed by ^53^). POA circuits that are activated by warming up can drive both sleep and cooling ^38^. However, Tb and physical activity were not changed by chemogenetic inhibition of POA^Hrh1^ neurons during the light/resting phase indicating that POA^Hrh1^ neurons are not required for this physiology.

Nest building is a component of sleep preparation ^53, 54^, maternal behavior ^39, 55^, and defense from cold ^19^. It is regulated by the POA ^7, 55, 56^ ^57^ and LH ^54^. POA neurons associated with nest building include PACAP-expressing (warm defense) and Brs3/Calcr-expressing (parenting) neurons. Nest building by individually housed male mice was decreased with chemogenetic inhibition of POA^Hrh1^ neurons, suggesting that parenting is not the driving behavior. Interestingly, activation of PFC^Sst^→LPO terminals increased nesting and Tb, but not sleep. The relevant LPO neurons are likely GABAergic and express *Meis2* and *Arpp21* ^56^, markers that correspond to *Hrh1*-expressing cluster i2. There was a strong Fos response in MnPO^Hrh1^ neurons to cold exposure. This could suggest that there are cold-responsive POA^Hrh1^ neurons that can drive nest building.

While Tb changes slightly lag the autonomic activity driving them, stimulation of POA^Hrh1^ neurons caused a very prolonged increase in Tb, longer than produced by similar activation of POA^Brs3^ neurons. We observed that the prolonged Tb increased was not due to prolonged Ca^2+^ activation of the POA^Hrh1^ neuron cell bodies. Activation of other POA populations can drive long-lasting reductions in Tb. For example, chemogenetic activation of Esr1a, Ep3r, and Trpm2-expressing populations produce hypothermia that extends far beyond the window of DREADD activation by CNO ^31, 33, 34^. These data suggest that multi-hour responses to short-term acute activation of Tb-controlling neurons is a general feature of Tb control, whether to increase or decrease Tb.

The Tb increase attenuated with daily CNO stimulation of the POA^Hrh1^ neurons. This is despite intact hM3Dq activation demonstrated by persistence of the physical activity increase. In other examples, daily CNO activation of hM3Dq on POA^LepR^ or DMH torpor-trapped neurons also showed a lack of attenuation, with persistently reduced Tb ^32, 58^.

Circadian Tb regulation is a ubiquitous feature of endotherms ^20^. We now identify a group of POA neurons contributing to the dark period Tb increase. Our data suggest that the POA^Hrh^ neurons contribute to maintaining Tb during periods of low activity and/or low Tb. Importantly, the POA^Hrh^ neurons do not affect dark phase physical activity. Besides the POA, other hypothalamic nuclei generally linked to circadian regulation include the SCN, Arc, and DMH ^31, 59, 60, 61^.

One of the major pre-autonomic output pathways of the SCN is to the subparaventricular zone (SPZ). Lesions of the dorsal SPZ, but not the PVH, specifically abolish Tb circadian rhythm, while keeping physical activity intact ^62^. The POA^Hrh1^ neurons receive synaptic input from both SCN and SPZ, possibly contributing to the circadian Tb regulation.

In conclusion, POA^Hrh1^ neurons participate in several thermoregulatory functions, including Tb regulation, physical activity, and nest building behavior. Our data suggest that these neurons control daily body temperature rhythm changes and cold-induced nesting behavior. Discrepancy between very strong Tb acute activation phenotype and the modest contribution to the dark phase Tb increase hints that POA^Hrh1^ neurons participate in unexplored thermogenic roles.

## METHODS

### Animals

Mouse studies were approved by the NIDDK Animal Care Use Committee (Animal Study Proposal K016-DEOB-23). Mice were on a 6 am – 6 pm lights on cycle at 22-24 °C (unless otherwise indicated). Mice had *ad libitum* access to chow (Teklad F6 RodentDiet 8664 or LabDiet cat# 5018; 3.05 kcal/g) and water, including during indirect calorimetry, chemo- and optogenetic telemetry and nesting experiments, but not during photometry, temperature preference and real-time place preference experiments. According to NIH guidelines, mice received twice daily status checks during surgery recovery and otherwise once. Mice were singly housed after telemetry probe and/or cranial implantation surgeries. Mice used were: Hrh1-Cre (generated in house; see below); B6.Cg-Hrh1^em1Rei^/J, (Hrh1^-/-^; Jax# 032615; RRID:IMSR_JAX:032615) ^63^; B6.Cg-Gt.(Rosa)26Sortm6(CAG-ZsGreen1)Hze/J, with Cre-dependent ZsGreen (Ai6; Jax# 007906; RRID:IMSR_JAX:007906) ^64^; Vglut2-FlpO (Jax# 030212; RRID:IMSR_JAX:030212); Vgat-FlpO (Jax# 029591; RRID:IMSR_JAX:029591); C57BL/6J; Brs3-T2A-FlpO (Jax#:040970; RRID:IMSR_JAX:040970 ^39^; Brs3-ires-Cre (Jax# 030540; RRID:IMSR_JAX:030540).

Male and female mice were used as indicated. We did not track the estrous cycle of female mice. For all experiments, mice were between 8 and 45 weeks of age. Mice from multiple litters were used for all experiments. Most Hrh1-Cre mice used were heterozygous for the Cre allele.

### Generation of Hrh1-Cre mice

CRISPR/Cas9 was used to cut at the Hrh1 stop codon in B6D2F1/J one-cell embryos, and a mouse with the correctly targeted insertion was identified and confirmed by sequencing. The targeting vector contained 3956 bp of C57BL/6 genomic sequence upstream of the Hrh1 stop codon, a T2A peptide sequence (54 bp), Cre recombinase (1083 bp), and 4191 bp of downstream C57BL/6 sequence. The T2A sequence adds EGRGSLLTCGDVEENPG to the C-terminus of Hrh1 and a proline to the N-terminus of Cre ^65^. PCR genotyping for Hrh1-Cre (496 bp) vs wildtype Hrh1 (369 bp) alleles uses x619 (Hrh1 forward, 5’-GCCAAGCAGTTGGGTTGTAT), x661 (Cre reverse, 5’-AATCGCGAACATCTTCAGGT), and x622 (Hrh1 reverse, 5’-ATGCCTTCACACTGCTCA). The founder was bred to C57BL/6J mice for >10 generations. Of note, the Hrh1 and Rosa26 locus are 1.3 Mb apart, resulting in low feasibility to bread double transgenic mice with a Rosa26 locus reporter.

### Hrh1^-/-^ mice

Hrh1^-/-^ and C57BL/6J (wt) mice were implanted with a Tb telemetry probe (Emitter; see below). After recovery, mice were acclimated to the experimentation room for 2 days. 4 to 6 days Tb and physical activity data collection, spread over two sessions, were averaged to 24h (ZT6 to ZT6) periods. In one of the two sessions, mice received a veh and a PEA (H_1_ agonist) injection at ZT4. PEA causes transient (1 h) of < 2 °C hypothermia in wt but not in Hrh1^-/-^ mice ^63^ The two hours around injection were removed from analysis.

### Stereotaxic injections

Mice were anesthetized with 0.5-1.5 % isoflurane, scalp hair was removed, and mice were placed in the stereotaxic apparatus (Digital Just for Mouse Stereotaxic Instrument, Stoelting) and eye ointment was applied (Puralube, Dechra). POA injections were all aimed at the anteroventral POA/MnPO. Viral vectors were injected with pulled glass pipette (20-40 µm tip diameter; 0.275 mm ID, 1 mm OD, Wilmad Lab Glass) at a rate of 50 nl per minute (visually controlled) with an air MAP system regulator (Grass Technologies, Model S48 Stimulator), after which the pipette was kept in place for 5 minutes. Mice received slow-release buprenorphine injections (0.1 mg/kg, s.c.; once) and ibuprofen in the drinking water for 5 days.

### Immunohistochemistry

Tissue preparation: mice were anesthetized with chloral hydrate (500/mg/kg in saline; i.p.), and perfused transcardially with ice cold saline followed by 10 % neutral buffered formalin. Brains were removed and post-fixed overnight (room temperature) and incubated in 30% sucrose (in 0.1M PBS; 4 °C) until sectioned (coronally; 50 µm) on a sliding microtome (SM2010 R, Leica) and collected in three series in 0.1 M PBS. Sections were washed, 3 x 10 minutes, in 0.1M PBS. Sections were incubated for 1 hour in 0.3% Triton X-100 with 3% normal goat serum in 0.1M PBS (PBS-T-NGS) and then moved to incubate overnight (room temperature) on a horizontal shaker in primary antibody. We used the following antibodies and dilutions in PBS-T-NGS: rabbit anti-DsRed at 1:2000 (632496, Clontech; RRID:AB_10013483) for mCherry; chicken anti-GFP at 1:1000 (13970, Abcam; RRID:AB_300798) for EYFP, GCaMP or GFP; guinea pig anti-Fos at 1:1000 (226-004, Synaptic Systems; RRID:AB_2619946). Sections were washed, incubated (2 h) with secondary antibody (1:500; goat-anti-rabbit Alexa-555 and/or Alexa-488 goat-anti-chicken and/or Alexa-594 anti-guinea pig; respective RRIDs, AB_2535849, AB_2534096, AB_2534120; Thermo Fisher) in PBS-T-NGS. After incubation with secondary antibody, sections were washed, mounted and cover slipped with Prolong mounting medium containing DAPI (Thermo Fisher).

### Viral vectors

We used the following viruses: AAV9-ef1a-DIO-ChR2(H134R)-EYFP (gift from K. Deisseroth; Addgene viral prep 20298-AAV9; http://n2t.net/addgene:20298 ; RRID:Addgene_20298); AAV1-CAG-FLEX-egfp (gift from Hongkui Zeng; Addgene viral prep # 51502-AAV1; ; http://n2t.net/addgene:51502 ; RRID:Addgene_51502; ^66^) AAV8-Ef1a-DIO-synaptophysin-mCherry (Virovek); AAV8-hSyn-DIO-hM4D(Gi)-mCherry (gift from B. Roth; Addgene viral prep # 44362-AAV8; http://n2t.net/addgene:44362; RRID:Addgene_44362), AAV8-hSyn-DIO-hM3D(Gq)-mCherry (gift from B. Roth; Addgene viral prep 44361-AAV8; http://n2t.net/addgene:44361 ; RRID:Addgene_44361) ^67^; AAV8-syn-DIO-mCherry (Addgene viral prep # 50459-AAV8; http://n2t.net/addgene:50459 ; RRID:Addgene_50459); AAV8-Ef1a-fDIO-GCaMP6s (gift from Rylan Larsen; Addgene viral prep # 105714-AAV8; http://n2t.net/addgene:105714 ; RRID:Addgene_105714); AAV8-CreOn/FlpOn-ChR2-EYFP (gift from Karl Deisseroth; Addgene viral prep # 55645-AAV8; http://n2t.net/addgene:55645 ; RRID:Addgene_55645); AAV8-CAG-FLEX-RabiesG(Y733F) (Stanford University Vector Core) EnvA-G-Deleted-Rabies-Egfp (Gene Transfer, Targeting and Therapeutics Core at Salk Institute); AAV8-Ef1a-DIO-synaptophysin-mCherry (Virovek, Inc.); AAV8-Ef1a-FLEX-TVA-mCherry (UNC Viral Vector core); AAV-hSyn-DIO-jGCaMP8s-P2A-ChrimsonR-ST (gift from Mark Histed (Addgene viral prep # 174007-AAV9 ; http://n2t.net/addgene:174007 ; RRID:Addgene_174007; ^68^; AAV-hSyn-DIO-HA-hM3D(Gq)-IRES-mCitrine (gift from Bryan Roth; Addgene viral prep # 50454-AAV8; http://n2t.net/addgene:50454 ; RRID:Addgene_50454)

### Chemogenetics

Hrh1-Cre mice were injected bilaterally with 25 nl of AAV8-hSyn-DIO-hM3D(Gq)-mCherry, 50 nl of AAV8-hSyn-DIO-hM4D(Gi)-mCherry or, as controls in the nesting experiments, 50 nl of AAV8-syn-DIO-mCherry in anteroventral POA (AP: 0.35; ML: 0.3; DV: -5.25 from bregma). DREADD expression was verified with immunohistochemistry for mCherry. All mice included in analyses had uni- or bilateral POA expression, with varying expression in POA subregions. Notably, some mice in the hM3Dq group lacked expression in the VMPO, but all had at uni- or bilateral expression in the MnPO or MPA. Most mice in the hM4Di group had bilateral MnPO (∼80%), MPA (95%) and VMPO (95%) expression. Both groups had some mice with expression outside of the POA (septum, BNST).

Experiments were performed as described below and in legends. All experiments were set up as randomized cross-over trials (vehicle vs CNO), unless otherwise specified. Mice received i.p. injections of 1.0 mg/kg CNO. Experiments were performed at 22 °C, unless indicated otherwise.

After two days acclimation to the experimentation room, Tb was recorded as a baseline, mice were randomized into vehicle or CNO groups and received injections for 5 days. After 4 days with no injection, mice received a recovery day injection on day 10. To calculate ΔTb the baseline Tb was subtracted from the daily Tb.

For cold/warm exposure Fos experiments, during the light phase, mice were habituated in a temperature-controlled chamber for two days prior to experiments in home cages on telemetry receiver boards. Ambient temperature was changed 30 minutes before dosing. Cold exposure lasted 4 hours at 5 °C and warm exposure 4 hours at 32 °C.

For LPS and Poly(I:C) experiments were conducted as a 2×2 factorial design with four treatment groups. Prior to this experiment, mice had been handled and injected with veh or CNO but were LPS or Poly(I:C) naïve.

To test nest building behavior, mice were habituated in a temperature-controlled chamber for two days prior to experiments. Existing nesting material (crinkle paper) was removed one hour prior to dosing and ambient temperature was adjusted as reported at that time. Five minutes after dosing, 6 g of crinkle paper was sprinkled on the cage floor ^69^. Four hours later, crinkle paper in nest and portion not in nest was weighed. Fraction used in nest was calculated. Notably, total mass of crinkle paper increased slightly during experiments, likely due to urine and/or humidity.

For the floor temperature preference test, two temperature choice tests were done by using the thermal place preference test (TPPT) instrument (Bioseb, Chaville, France). We first tested C57BL/6J male mice under several two temperature combinations for 15 min to find the temperatures in which mice stay in either temperature zone for roughly 50% of the total time, found that the two temperatures of 25°C vs 38°C was the optimal combination, and subsequent experiments were run using these temperatures. For each run, two mice were placed in a plexiglass chamber separated by a black plastic plate in the middle, with two adjacent thermal surfaces both with an accuracy of ±0.1°C. Mouse movements were recorded with a video tracking system for 15 min. Two to three hours later, mice were tested in the opposite lane, with the plate temperatures reversed. Time in 25°C (vs 38°C) zone was calculated by the software. An average of the two 15 min test periods were calculated and used for graphing. Female mice prefer warmer temperatures in this assay^70^.

### Body temperature telemetry

For surgery, animals were anesthetized with 0.5-1.5 % isoflurane, the abdominal hair shaved and E-Mitters (Starr Life Sciences) were implanted in the intraperitoneal cavity ^71^. Emitters were sutured to the abdominal wall, however at euthanasia they were rarely still attached. To account for slight differences that changed emitter positions may cause, Tb quantification is represented as ΔTb, unless otherwise specified.

For the telemetry experiments, mice were housed in their home cages on ER4000 energizer/ Receivers (Starr Life Sciences) at least two days before experiments to acclimate to the temperature-controlled chamber or dedicated telemetry room. The Tb and physical activity were continuously measured as 1-min means by VitalView software (Starr Life Sciences). Experiments were performed at 22 °C, unless indicated otherwise. Physical activity is measured in counts (arbitrary units).

### Indirect calorimetry

An Oxymax/CLAMS (Columbus Instruments) was used for continuous monitoring of energy expenditure, food intake, water, respiratory exchange ratio, physical activity (beam breaks), and body temperature (from Emitters). Mice were acclimated to the chambers two days prior to start of experiments. Experiments were performed at 22 °C. Each cage was sampled every 260 seconds.

The physical activity energy expenditure (PAEE) was calculated as described ^27^ and subtracted from the TEE (TEE-PAEE). The calculated PAEE is an upper bound, since it does not consider the compensatory reduction in CIT that occurs below thermoneutrality.

### Optogenetics

Hrh1-Cre mice were injected with 50 nl of AAV9-Ef1a-DIO-ChR2(H134R)-EYFP or, as controls, with AAV1-CAG-FLEX-EGFP in the right anteroventral POA (AP: 0.35; ML: 0.3; DV: -5.25 from bregma). For intersectional experiments, Hrh1-Cre;Vgat-FlpO or Hrh-Cre;Vglut2-FlpO mice, or Hrh1-Cre as controls, were injected with 100 nl of AAV8-CreOn/FlpOn-ChR2-EYFP at the same coordinates. Optical fibers were made in house (200 mm diameter core; NA 0.22; Nufern) were glued to ceramic zirconia ferrules (230 mm bore; 1.25 OD diameter; Precision Fiber Products) or premade (CFMLC12U; Thorlabs) and cleaved at desired length. Optic fibers were implanted unilaterally over the POA (AP: 0.35;ML: 0.3; DV: -4.5, mm from bregma), DMH (AP: -1.85; ML: 0.3; DV: -4.5, mm from bregma), Arc (AP: -1.6; ML: 0.2; DV: -5.5, mm from bregma) or VTM (AP: -5.7; ML: 1.0; DV; -5.0, mm from bregma). Fibers were attached to skull with C&B Metabond Quick Cement and dental acrylic. After completion of experiments, fiber placement and ChR2 expression were assessed. Animals without ChR2 expression or incorrect placement of optic fibers were excluded from analysis.

Fiber optic cables (200 mm diameter, NA 0.22, 0.5mlong, ThorLabs) were connected to the implanted fiber optic cannulas with zirconia sleeves (Precision Fiber Products) and coupled via a fiber optic rotary joint (Doric Lenses). We adjusted the light power of the laser (473nm; Laserglow or OptoEngine) at the end of the fiberoptic to ∼10 mW (measured with a fiberoptic power meter; PM20A; ThorLabs). The estimated light power at tip of implanted fiber between 3 and 6mW/mm^2^ (calculated according https://web.stanford.edu/group/dlab/cgi-bin/graph/chart.php). This is an upper limit due to possible light loss between the fiber optic cable and the implanted optic fiber. Light pulses were controlled by a programmable waveform generator (Arduino). 10 ms pulses were delivered at 20 Hz for 1 second, followed by 3 seconds off, except for real-time place preference experiments in which pulse trains were on for 1 second, followed by 2 seconds off. Frequencies were adjusted accordingly in the frequency response experiment.

For Tb telemetry experiments, mice were acclimated to the attached fiber patch cord for at least two days immediately before experiments in their home cage and typically were not handled on the day of the experiment. Photostimulation was done during the light phase or dark phase as indicated.

We used quantitative thermal infrared imaging of shaved skin temperatures in the interscapular and dorsal lumbar regions were used as measures of BAT and core body temperature, respectively ^25, 72, 73,74^. One day before experiments, the interscapular region and a 2 x 2 cm midline area 2 cm above the tail were shaved under isoflurane anesthesia. After overnight housing in their home cage with the optical fiber attached, mice were placed in a 20 x 20 cm enclosure with bedding, 170 cm below the infrared (IR) camera (T650sc, FLIR Systems, Wilsonville, OR), and acclimated for 3 h. Mice then underwent 2 cycles of 20 min baseline, 20 min laser on, and 40 min post-stimulation, with IR images collected continuously using ResearchIR software (FLIR Systems, Wilsonville, OR; 7.5 frames per second; IR emissivity set to 0.97 ^25^. Maximum interscapular and lumbar temperatures were determined by a blinded observer every minute using FLIR Tools (FLIR Systems, Wilsonville, OR). The ambient temperature was 25-26 °C.

For real time place preference, the day before the experiments, mice were connected to fiber optic cable to acclimate and placed in a two-chamber custom-made enclosure: white plexiglass walls and floor (50 × 26 × 30 cm) with a middle partition with opening in the center, allowing the mice to move between sides. On the day of the ‘no laser’ (control) experiment, mice were connected and video recorded for 20 minutes. The next day, one side was paired with optogenetic stimulation and the other was not. Laser was controlled with Arduino and was programmed with Ethovision to deliver pulses only when the mouse was one side of the chamber. Laser side was switched after each run. Movement tracking analysis was performed with Ethovision XT 14 software (Noldus). Experiments were done during the light phase.

For food intake optogenetic experiments, mice were housed in their home cage and connected to the fiber optic two days prior to experiments. Food was removed and mice were given a fresh cage and allowed to habituate for an hour. For food intake during stimulation experiments, a food pellet was weighed, delivered to the cage and optogenetic stimulation was started. After one hour the food pellet was weighed. For post stimulation food intake experiments, mice underwent one hour of optogenetic stimulation in a fresh cage with no food and then received a food pellet that was weighed after one hour.

For the frequency dose response experiment, mice were housed in their home cage and connected to the fiber optic two days prior to experiments. Ten minute stimulation at each frequency was done on two separate days one week apart (randomized frequency; order varied between days) in three mice, 3 Hz only has one repeat due to technical reasons. ΔTb is Tb at 10 minutes after laser start minus Tb at 0.

For the brown adipose tissue denervation, we cut sympathetic nerves of the interscapular brown adipose tissue as described^75^. In brief, POA^Hrh1^::ChR2 mice were acclimated in their home cage to 30 °C for 5 days prior to pre-denervation optogenetic testing (Pre). Mice were anesthetized with isoflurane, the surgical area was shaved, and a ∼1.5 cm skin incision was made. The fat pads were detached from underlying muscle and raised to expose the BAT ventral surface. On both sides, the five nerves (some are in bundles) were identified, dissociated from the pad and cut in two places, removing a section of nerve. This removes both sympathetic and sensory innervation. In sham animals the BAT and nerves were handled similarly but not cut. The fat pads were rinsed with saline and the incision closed with dissolvable sutures. Mice recovered for 4 days at 30 °C to reduce cold stress before the post-denervation optogenetics (Post). No change in baseline Tb was observed in the denervation group.

### Single-cell RNA sequence database analysis

The scRNA seq count matrix for GSE113576 ^30^ was analyzed with R (v. 4.0.3) using Seurat (v.3.2.2) as described before ^4^. Out of 18,401 neuronal cells, we re-clustered the 1701 neurons with R1 UMI corresponding to Hrh1 (PC: 20, resolution: 0.6). Clusters labeled beginning with ‘i’ represent inhibitory clusters enriched with Vgat, while those labeled with ‘e’ represent an excitatory cluster enriched with Vglut2. An ‘apoptotic’ cluster was enriched for several mitochondrial genes, suggesting cell death, and left out of the graphs.

### Drugs

Clozapine-N-oxide (CNO; Sigma C0823) stock was made by dissolving in sterile 0.9% saline at 0.5 mg/ml and frozen. On the experimental day it was diluted and administered at 1 mg/kg i.p. Lipopolysaccharide from Salmonella enterica serotype typhimurium (LPS; Sigma L6511) was dissolved in sterile saline and administered at 50 mg/kg i.p. Poly(I:C) (Tocris 4287) was dissolved in 0.9% saline and administered at 10 mg/kg i.p. Pyrilamine maleate (Sigma SML3651) was dissolved in 0.9% saline and administered at 10 mg/kg i.p. 2-Pyridylethylamine (PEA, Tocris 2478) was dissolved in ddH_2_O, frozen at 10 mM and used at 100 µM.

### Neuronal tracing

For anterograde tracing experiments, Hrh1-Cre mice were injected with <10 nl of AAV8-Ef1a-DIO-synaptophysin-mCherry in the MnPO/anteroventral POA. Mice were euthanized 6 weeks after injection and brains processed for immunohistochemistry (see above). All 6 mice had expression in cell bodies limited to the anterior POA had consistent axonal projections to described areas. For each target region, the density of synaptophysin-mCherry–positive puncta and fibers was scored by eye on a 4-point ordinal scale: 1 = very sparse labeling, 2 = sparse labeling, 3 = moderate labeling, and 4 = dense labeling (numerous puncta/fibers) throughout the region.

For monosynaptic rabies tracing, we injected 100 nL of a 1:1 mixture of AAV8-Ef1a-Flex-TVA-mCherry and AAV8-CAG-FLEX-RabiesG in the MnPO/anteroventral POA of Hrh1-Cre mice. 4 weeks later, we injected 150 nL EnvA-G-deleted-Rabies-GFP in the DMH or POA. 10 days after rabies virus injection, mice were euthanized and their brains were processed for IHC (see above). Brain regions were defined according to the *Mouse Brain Atlas*.^76^ Neurons were counted as starter neurons when mCherry fluorescence was detected in GFP+ neurons.

For POA^BRS3^→DMH collateral tracing, AAV-DIO-TVA-mCherry was injected in the right MnPO/anteroventral POA of Hrh1-Cre mice (50 nl). After >4 weeks retrograde G-deleted-Rabies-GFP (100 nl) was injected in the right DMH. Mice were perfused 10d after last injection and brains processed for immunohistochemistry (see above) for GFP and mCherry.

For anterograde tracing from POA^Hrh1;Vgat^ or POA^Hrh1;Vglut2^ neurons, brains from POA^Hrh1;Vgat^ or POA^Hrh1;Vglut2^ mice in the intersectional optogenetics experiments were processed for IHC (see above). DMH, Arc and VTM were analyzed for projections (n=3 for each group).

### Fos experiments

For 4 °C vs 30 °C ambient temperature experiments, mice used for hM4Di experiments were habituated for two days to the thermal chamber in their home cages. Ambient temperature was changed to 30 °C (or 4 °C) at ∼10 am. After 3 hours, mice were anesthetized and transcardially perfused. 1-2 slices were counted for each brain region (see Fig 8a for delineation of regions). Counting was done blinded to group. There was no difference in number of neurons per region between treatments

For ZT8 vs ZT14 experiments, mice used for hM3Dq experiments were habituated to an alternative light/dark schedule (3 pm lights off; 3 am lights on) for a week in a thermal chamber (22 °C), to minimize perturbance. On day 8 at ZT8 and day 9 at ZT14, mice (3 males, 3 females in each group) were anesthetized and transcardially perfused. 1-2 slices were counted for each brain region (see Fig 8a for delineation of regions). VMPO did not consistently contain sufficient hM4Di-mCherry-expressing neurons (range: 0 to 114)and was omitted form analysis. Counting was done blinded to group. There was no difference in number of neurons per region between treatments.

### Western blot

Mice were euthanized at ∼1 pm and interscapular brown adipose tissue (BAT) was dissected and stored at -80 °C. Protein from BAT was extracted with RIPA buffer, quantified (Pierce BCA Protein Assay Kit, Thermo Scientific, Rockford, IL), separated in 4–20% SDS-PAGE gels (17 μg/lane), transferred to PVDF membrane, and probed with anti-TH (1:5000, Abcam, ab137869), anti-UCP1 (1:5000, Sigma-Aldrich, U6382), or anti-α-Tubulin (1:5,000, Sigma–Aldrich, T6074). Signals were detected with SuperSignal West Pico Chemiluminescent Substrate (Thermo Scientific), and images were scanned with ChemiDoc Touch Imaging System (BIO-RAD).

### BAT histology

BAT tissues were dissected from mice, fixed in 10% formalin, embedded in paraffin, sectioned, stained with hematoxylin and eosin, and examined by light microscopy.

### Photometry

Hrh1-Cre or Brs3-Cre male mice were injected with 50 nl of AAV9-syn-DIO-jGCaMP8s-P2A-ChrimsonR-ST or, as controls, with AAV8-syn-DIO-HA-hM3Dq-mCitrine, in the right anteroventral POA (AP: 0.35; ML: 0.3; DV: -5.25 from bregma). Optical fibers (fiber: core = 400 μm; 0.48 NA; M3 thread titanium receptacle; Doric Lenses) were implanted over the right POA (AP: 0.35;ML: 0.3; DV: -5.0, mm from bregma). Fiber and a custom-made titanium head post (H.E. Parmer) were attached to skull with C&B Metabond Quick Cement (Parkell) and dental cement (Stoelting) for optogenetic and pyrilamine experiments.^77^ Mice were allowed to recover at least three weeks before experiments.

For optogenetic stimulation (head-fixed) and pyrilamine injection (freely moving) experiments, mice were habituated to head-fixation and with the ability to locomote on a 3D-printed circular treadmill ^77^on the two days preceding recording day. Fiber optic cables (Low-Autofluorescence Mono Fiber-optic Patch Cord; MFP_400/430/-0.57_2m_FCM-CM3_LAF) were connected to implanted optic fiber. Excitation and emission light was passed through a fluorescence minicube allow for 405 nm and 465 nm excitation (5 port; Doric Lenses; FMC5_IE(400-410)_E(460-490)_F(500-540)_O(580-680)_S). Excitation light (∼100 μW) was provided by a 405 nm or 465 nm LED (Plexon). Excitation LEDs were modulated as interleaved pulses controlled by the Nanosec photometry-behavioral system that was initially developed in a previous study ^78^ (https://github.com/xzhang03/NidaqGUI). Emission light was collected by a femtowatt photoreceiver (Newport 2151) and digitized at 2.5 kHz sampling rate (PCIe-6321; National Instruments). Data acquisition was controlled using a modified script in MATLAB (MathWorks) based on our previously developed script^78^. Arduino-controlled optogenetic stimulation with a 635 nm laser (Shanghai Laser) delivered 10 ms pulses at 20 Hz for 1 second, followed by 3 seconds off. During the first five minutes of recording heavy bleaching occurred and data were discarded in all recordings. Data analysis was conducted in MATLAB by first applying a least-squares linear fit to the 405 nm signal to align it to the 465 nm signal. The resulting fitted 405 nm signal was then used to normalize the 465 nm signal as follows: 465 nm signal − fitted 405 nm signal. The normalized fluorescence signal (F) was then z scored: (F- (mean [F_baseline_])/SD[F_baseline_]. F_baseline_ is -5 to 0 in the optogenetic stimulation and -10 to 0 in the pyrilamine experiment. Quantification in the optogenetic experiment: post early is the Δz score (10 to 11) – (-1 to 0) minutes and post late is Δz score (25 to 30) – (-5 to 0) minutes. Quantification for the pyrilamine experiment is: Δz score (-10 to -1) – (11 to 20) minutes, avoiding handling artifacts. The optogenetics experiment was repeated and data averaged.

For homecage ZT8-14 recording, a telemetry probe for Tb and physical activity was implanted in the intraperitoneal cavity (see above). Mice were kept on an alternative light/dark schedule (3 pm lights off; 3 am lights on) for at least a week before experiments. Mice were acclimated to the attached fiber patch cord for at least two days immediately before experiments in their home cage and were not handled on the day of the experiment. Photometry recordings were performed with Tucker-Davies Technologies system as previously described ^79^. The photometry and telemetry recordings were started a few minutes before ZT8. Some mice received a brief >1 s optogenetic pulse prior to ZT8 and after ZT14 to confirm that the GCaMP signal was preserved throughout the recording. During the first few minutes, heavy bleaching occurred and data were discarded. The calcium fluorescence signal was calculated as follows by calculating ΔF/F as (470-fitted405)/fitted405. The ΔF/F was then used to calculated z-scores (with baseline being the entire ZT8-14 period): ((ΔF/F) – (mean [F_baseline_]))) / SD[F_baseline_].

### Slice 2-photon imaging

Hrh1-Cre mice were injected with 50 nl of AAV9-syn-DIO-jGCaMP8s-P2A-ChrimsonR-ST bilaterally in the anteroventral POA (AP: 0.35; ML: 0.3; DV: -5.25 from bregma). After >4 weeks, as previously described ^80^ mice were anesthetized with isoflurane and decapitated. The brain was removed and immediately immersed in ice-cold, choline-based cutting solution (92 mM choline chloride, 10 mM HEPES, 2.5 mM KCl, 1.25 mM NaH2PO4, 30 mM NaHCO3, 25 mM glucose, 10 mM MgSO4, 0.5 mM CaCl2, 2 mM thiourea, 5 mM sodium ascorbate, 3 mM sodium pyruvate, pH 7.4) and sliced (300 µm) on a vibratome (Campden 7000smz-2). Slices were transferred to a warmed (36°C) choline cutting solution for 15 minutes, followed by continued recovery in 36°C oxygenated (95% O2/ 5% CO2) artificial cerebral spinal fluid (ACSF; 126 mM NaCl, 21.4 mM NaHCO3, 2.5 mM KCl, 1.2 mM NaHPO4, 1.2 mM MgSO4, 2.4 mM CaCl2, 10 mM Glucose) for 30 minutes. After slices were then kept at room temperature until used.

A single slice was transferred to a recording chamber (RC-26G; Warner Instruments) perfused with oxygenated ACSF at room temperature at a flow rate of approximately 2 mL/min. For imaging, a two-photon microscope (Olympus FVMPE-RS) equipped with an Insight laser was used at an excitation wavelength of 920 nm and resonant scanning was used to acquire 30 fps with a 20x 1.0 NA water-immersion objective (Olympus). After 10 minutes of baseline recording, we optogenetically stimulated with a 620 nm LED (1 mW/mm2, Luxeon Star LEDs) driven by an Arduino-controlled driver (Luxeon Star LEDs) below the chamber for 10 minutes (10 ms pulses at 20 Hz, 1 second on, 3 seconds off). After 10 minutes of recovery, PEA was added to the perfusion ACSF (100 µM). Image acquisition was performed using Olympus Fluoview software.^80^

Image registration for two-photon calcium images of brain slices was corrected for minor XYZ drift using Fast4Dreg (ImageJ v1.54f) and excluded if drift remained following this registration. ROIs generated from Cellpose (https://github.com/mouseland/cellpose) were then imported into the ROI manager and the mean intensity of each ROI was measured over time.

The first 5 minutes of the recordings (-10 to -5 minutes) were discarded to allow stabilized baseline recording. The change in fluorescence was calculated by generating the average intensity projection of baseline florescence (F0) during the baseline period (-5 to 0 minutes). Using the image calculator function in FIJI, F0 was subtracted from the fluorescent time course F and then the resulting image was divided by F0:

### ΔF/F0 = (F-F0)/F0

Cells were designated as responsive if the change in fluorescence was greater than 3 standard deviations from the baseline period (-5 to 0 minutes). For the response to PEA the mean fluorescence from 23-25 minutes was subtracted by then mean from 17-19 minutes. Data was further visualized by average and by heatmap of all cells.

### Statistics

Within subject comparisons used paired Student’s T-Tests for two conditions and one-way ANOVA for more than 2 conditions. Data are presented as mean ± S.E.M, with error bars omitted in some graphs for visual clarity. Data analyses were performed using R, Microsoft Excel, and GraphPad Prism.

## Abbreviations neuroanatomy

3V: Third ventricle
4V: Fourth ventricle
aca: Anterior commissure
AcbSh: Nucleus accumbens, shell
AH: Anterior hypothalamic area/nucleus
AHA: Amygdalohippocampal area
Arc: Arcuate nucleus
aq: Cerebral aqueduct
AVPe: Anteroventral periventricular nucleus
BNST: Bed nucleus of the stria terminalis
CLA: Claustrum
DB: Diagonal band
DMH: Dorsomedial hypothalamus (nucleus)
EPD: Dorsal endopiriform nucleus
f: Fornix
GiA: Gigantocellular reticular nucleus, alpha part
LH: Lateral hypothalamus
LM: Lateral mammillary nucleus
LPB: Lateral parabrachial nucleus
LPO: Lateral preoptic area
LS: Lateral septum
LSD: Lateral septal nucleus, dorsal
ME: Median eminence
MeA: Medial amygdala
ML: Medial mammillary nucleus, lateral part
MM: Medial mammillary nucleus, medial part
MnPO: Median preoptic nucleus
MPA: Medial preoptic area
MS: Medial septum
MTu: Medial tuberal nucleus
OVLT/VOLT: Vascular organ of the lamina terminalis (organum vasculosum)
PAG: Periaqueductal gray
Pe: Periventricular nucleus
PFC: Prefrontal cortex
PH: Posterior hypothalamic area
PM: Premammillary nucleus
POA: Preoptic area
PV: Paraventricular thalamic nucleus
PVH: Paraventricular hypothalamic nucleus
RCh: Retrochiasmatic area
RIP: Raphe interpositus nucleus
RMg: Raphe magnus nucleus
RPa: Raphe pallidus
scp: Superior cerebellar peduncle
SCh/SCN: Suprachiasmatic nucleus
SFO: Subfornical organ
SHy: Septohypothalamic nucleus
SON: Supraoptic nucleus
SPa: Subparaventricular area
SPZ: Subparaventricular zone
StHy: Striohypothalamic nucleus
TMN: Tuberomammillary nucleus
VLPO: Ventrolateral preoptic nucleus
VMH: Ventromedial hypothalamus
VMPO: Ventromedial preoptic area
VTA: Ventral tegmental area
VTM: Ventral tuberomammillary nucleus

## DATA AVAILABILITY

All data reported in this paper will be made available in the Source Data file. Any additional information required to reanalyze the data reported in this paper is available from the lead contact upon request.

## CODE AVAILABILITY

This paper does not report original code.

## ACKNOWLEDGEMENTS

We thank for their assistance: Alice Franks, Yuning Huang, Naili Liu, Tamar Demby, Chengyu Liu (NHLBI Transgenic Core), Danielle Lafferty and Kaitlyn Hajdarovic. This research was supported by the Intramural Research Program (ZIA DK075057; ZIA DK075063; ZIA DK075064; ZIA DK075168; ZIA DK075087-07; ZIC DK070002) of the National Institute of Diabetes and Digestive and Kidney Diseases (NIDDK) within the National Institutes of Health (NIH). The contributions of the NIH author(s) are considered Works of the United States Government. The findings and conclusions presented in this paper are those of the authors and do not necessarily reflect the views of the NIH or the U.S. Department of Health and Human Services.

## AUTHOR CONTRIBUTIONS

R.A.P, with input from M.L.R., A.L. and M.J.K conceived the project and designed the experiments. R.A.P., A.V., A.I.G. and S.K. performed stereotaxic surgeries. R.A.P, O.G. and A.V. performed chemogenetic indirect calorimetry and Tb telemetry experiments. R.A.P and A.V. performed nest building behavior experiments. R.A.P. performed and analyzed fiber photometry experiments. D.K. performed and analyzed 2P experiments. R.A.P and A.V. performed optogenetics experiments. R.A.P., A.V., A.I.G. and S.K. performed histology and antibody staining. C.K.H. analyzed and prepared figures of scRNA-seq data. C.X. and M.L.R. generated Hrh1-Cre mouse line. C.X performed floor temperature assay and Western blot. R.A.P. prepared figures. R.A.P. wrote the manuscript, which all authors reviewed and edited.

---

**sFig 1.**
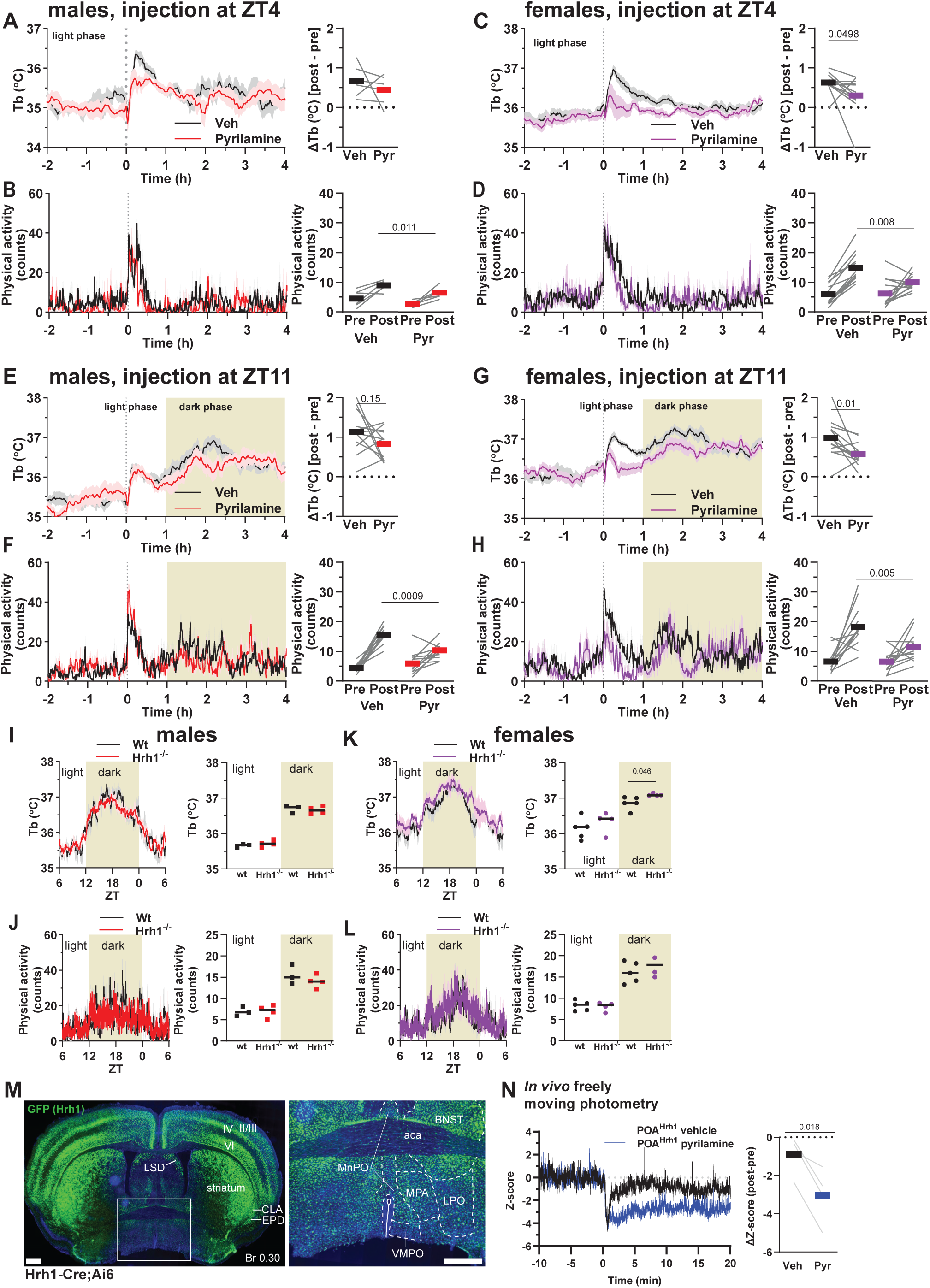
Histamine H_1_ receptor antagonist pyrilamine reduces Tb and POA^Hrh1^ neuronal activity. A-H) Tb and physical activity response to vehicle (Veh) or histamine H1 receptor antagonist pyrilamine. Right side, responses of individual mice (gray lines; mean indicated by bar). Injection is at time 0, data are mean ± SEM, males n = 6 – 12/group and females n = 13/group. In the light phase (A-D), pre is the mean of -120 to -30 minutes and post is the mean of 0 to 90 minutes. In the dark phase (E-H), pre is the mean of -120 to -30 minutes and post is the mean of 60 to 150 minutes. ΔTb is post minus pre. I-L) Tb and physical activity of wt and Hrh1-/- mice. Right side, responses of individual mice during 12 h light and dark intervals (data are mean ± SEM, male n = 3 – 4, and female n = 4 – 5). M) GFP reporter expression in hypothalamus of Hrh1-Cre;Ai6 mouse. Scale bar is 500 µm. N) In vivo freely behaving fiber photometry Ca2+ response of POAHrh1 neurons to a 10 mg/kg ip injection of pyrilamine. Right side, responses of individual mice, n = 4 males. A-H, N: P values are paired t-tests. I-L: P values are unpaired t-tests.

**sFig 2.**
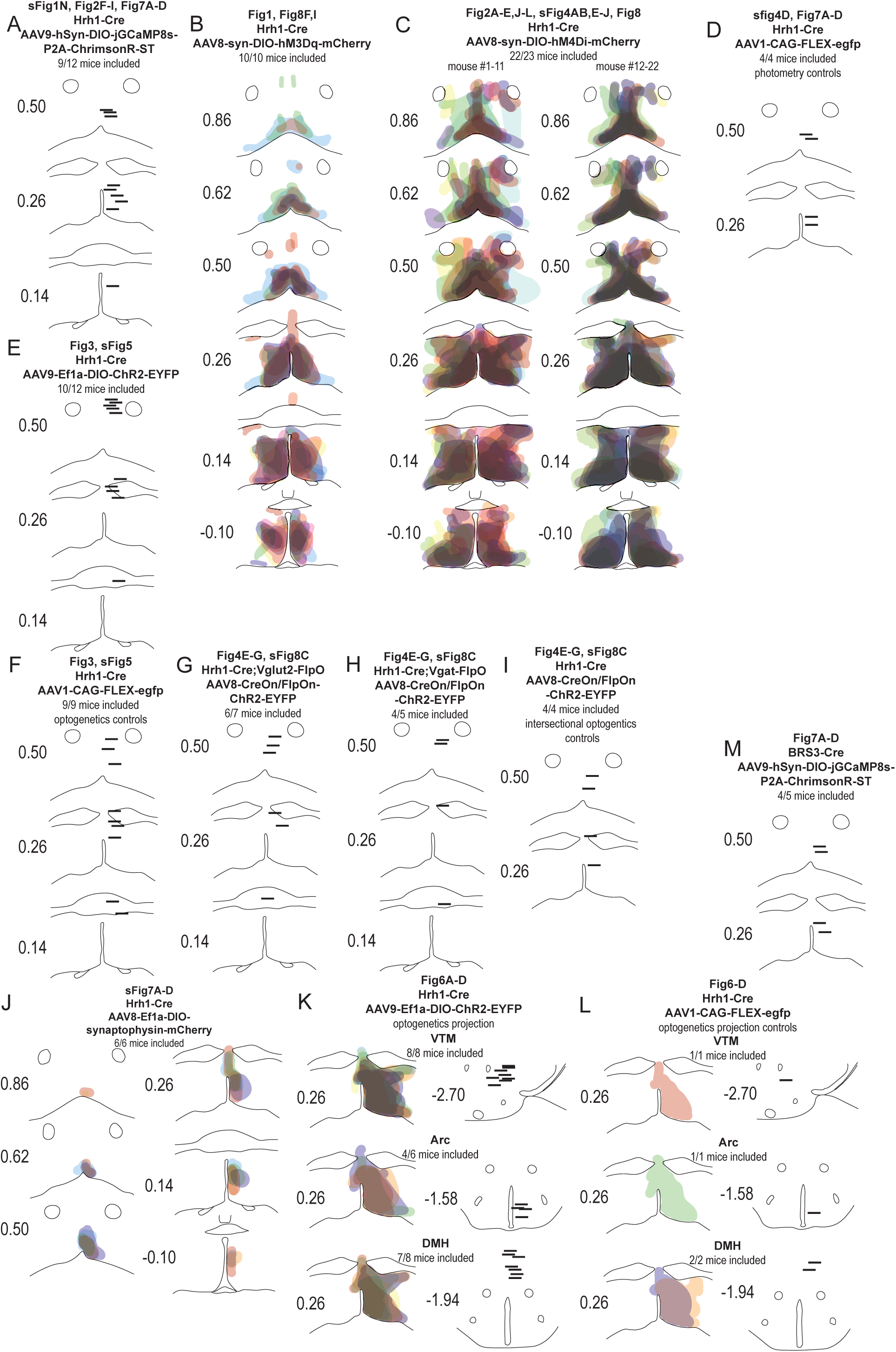
Histology summary of viral injections and optogenetic or photometry fiber placement. A-M) Each color represents one expression pattern in one mouse. Horizontal black lines represent the placement of optogenetic or photometry fibers. On the left of each schematic is the rostrocaudal position from Bregma in mm. Each schematic lists the number of mice with suitable targeting/number of the mice that underwent surgery.

**sFig 3.**
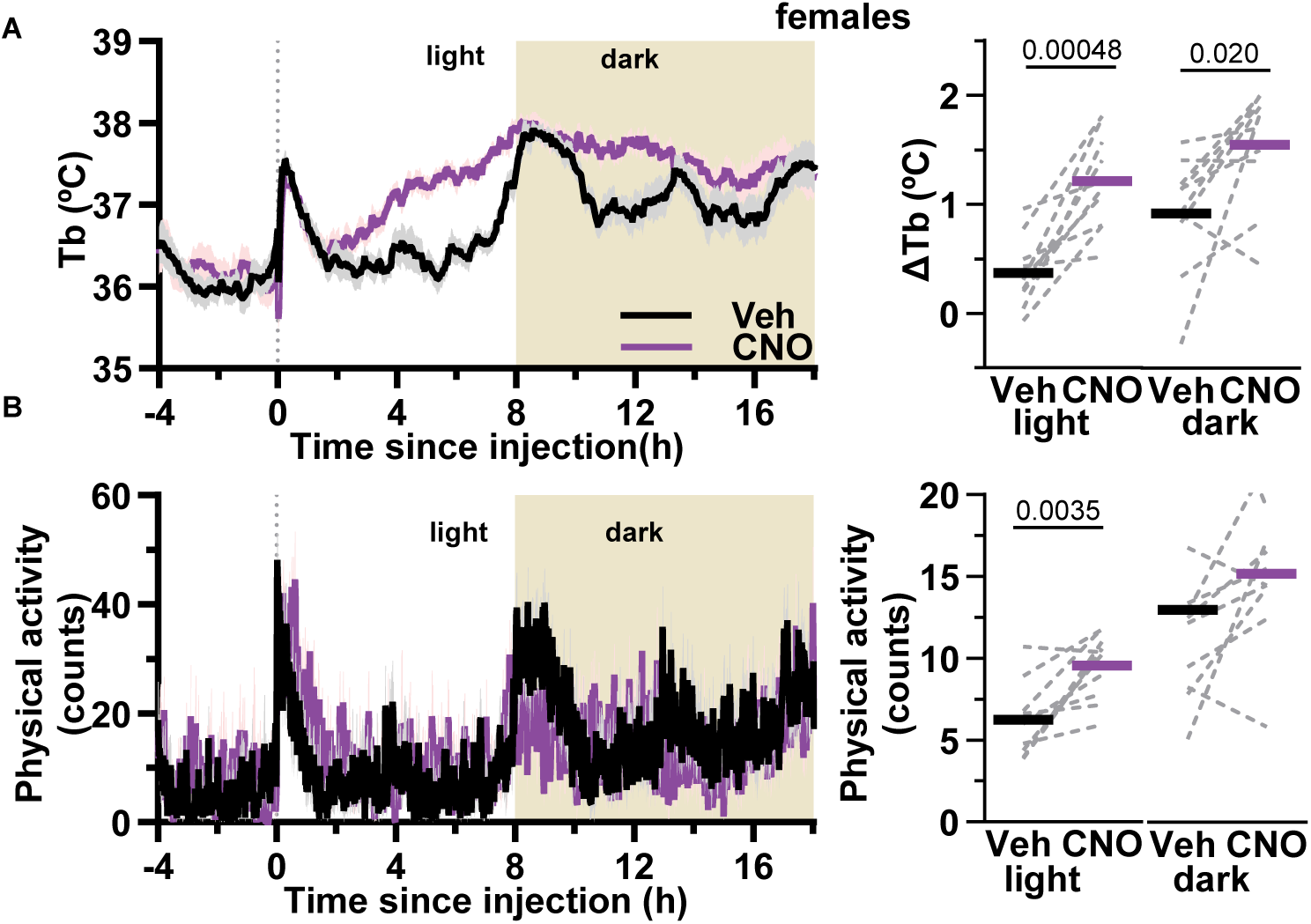
POA^Hrh1^ neuron activation in female mice. A) GFP reporter expression in hypothalamus of Hrh1-Cre;Ai6 mouse. Scale bar is 500 µm. B, C) Tb and physical activity response (mean ± SEM) to vehicle (Veh) or CNO injection (dotted line) in female mice (n = 10). Right side, quantification of mean response (bar) and individual mice (gray dotted lines) during light (A-D; Tb) and dark (E-H; 8 to 20 h) interval. ΔTb is interval minus baseline (-2.5 to -0.5 h) and physical activity is the mean during the interval. P values are paired t-tests.

**sFig 4.**
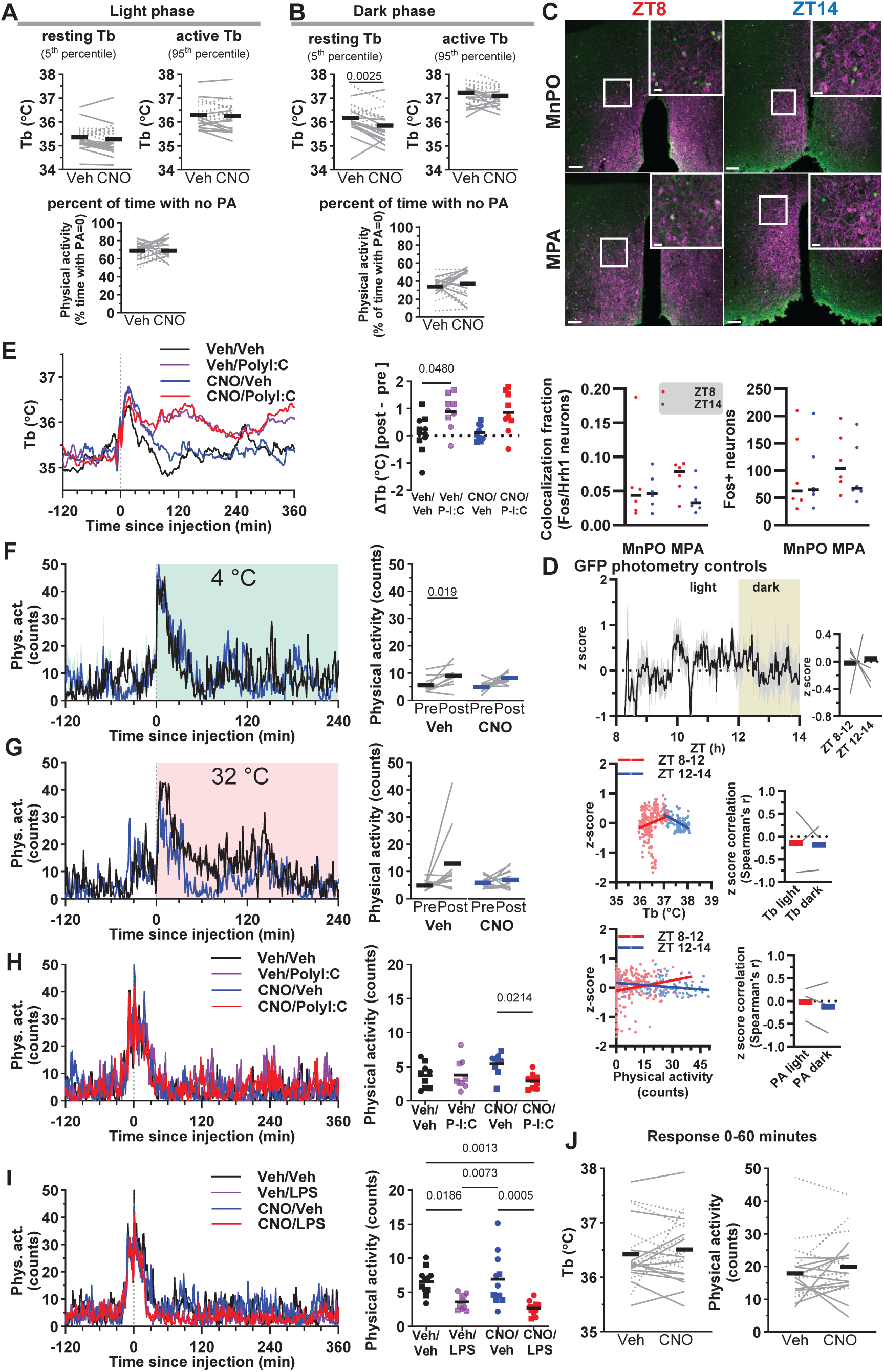
Chemogenetic inhibition of POA^Hrh1^ neurons reduces lowest 5^th^ percentile dark phase Tb increase values, but not physical activity. A,B) For light and dark phase, the resting (5^th^ percentile: the 5 % of lowest Tb values) and active (95^th^ percentile: the 5 % of highest Tb values) Tb were calculated. For physical activity, percent of time without physical activity was calculated. Analysis from 60 to 180 minutes. C) Top panels: examples of Fos expression at ZT8 and ZT14 in MnPO and MPA in Hrh1-expressing neurons. Scale bar is 50 µm; inset is 2x magnification. Quantification: (left) fraction of Hrh1 neurons that expresses Fos in MnPO (143 ± 23 mCherry neurons) and MPA (182 ± 23 mCherry neurons); (right) number of Fos neurons per nucleus in male and female mice (n = 6). D) Mean z-score of Ca^2+^ fiber photometry recording (±SEM) of POA^Hrh1^ neurons in control male mCitrine-expressing mice (n = 4) from ZT8 to 14 (top left). Quantification (top right): mean (bar) and individual mice (gray lines). Correlation of z-score with Tb (midle panels) and physical activity (lower panels); quantification (right): mean (bar) and individual mice (gray lines). E) Mean Tb responses to chemogenetic inhibition with PolyI:C treatment (left). Quantification (right): ΔTb, post (mean of 60 to 180 minutes) minus pre (mean of -150 to -30 minutes). F-I) Quantification physical activity (right; pre is -150 to -30 minutes and post is 60 to 180 minutes; mean (bar) and individual mice (gray lines or symbols). J) Tb and physical activity quantification during the first hour after ip injection of analysis of mean of three experiments: 1) Fig2 B,C, veh vs CNO; 2) Fig2 L, sFig 4I, Veh/Veh vs CNO/Veh; 3) veh vs 5mg/kg CNO. E-J) Mean (bar) and individual mice (males: uninterrupted lines or squares; females: dotted lines or circles). In E) and I) t=0 aligned with second dosing. P value: in A-B, D, F-G, J paired t-test; in C unpaired t-test; in E, H-I one-way ANOVA with Tukey’s multiple comparison test comparing all means to each other.

**sFig 5.**
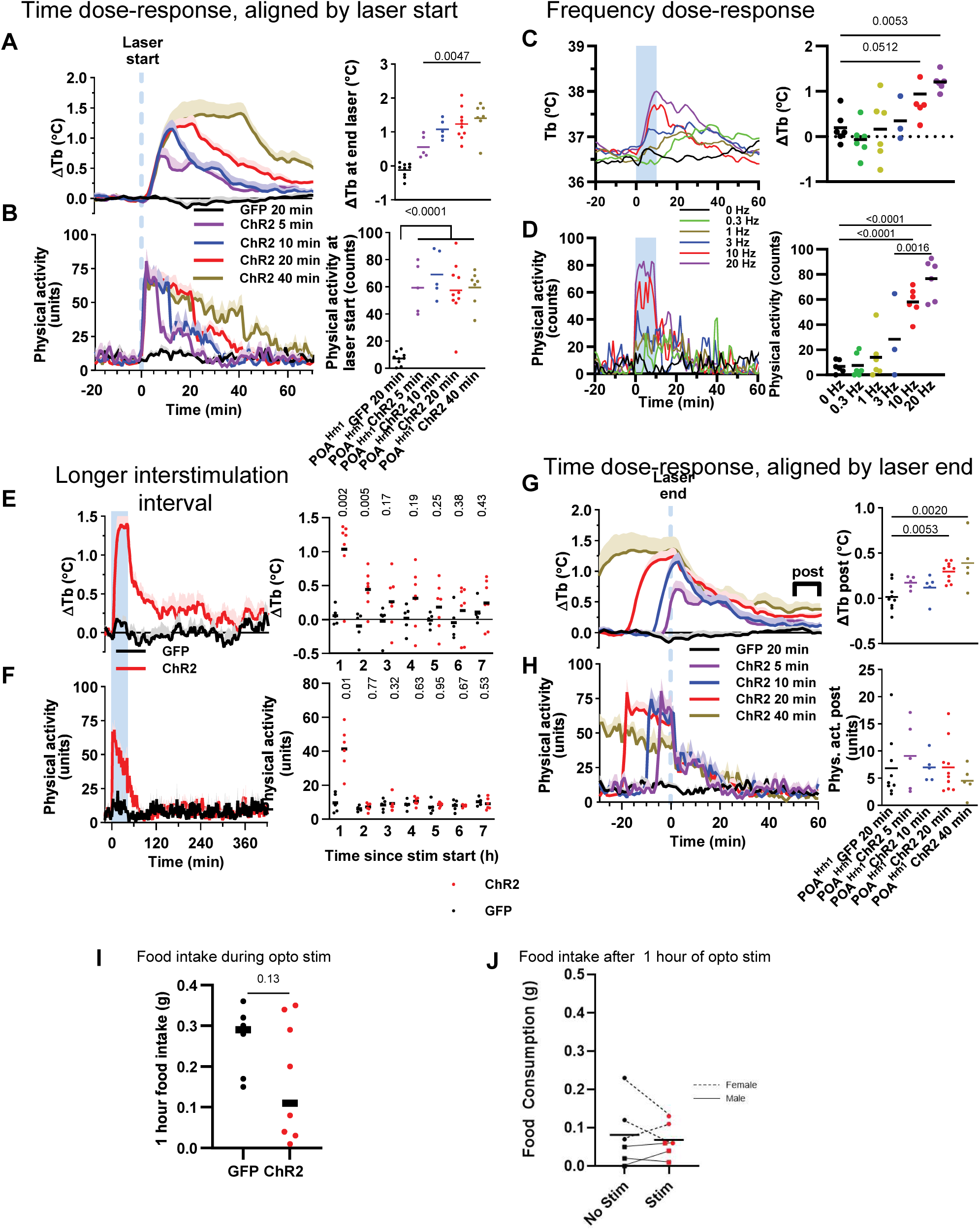
Body temperature, physical activity, and food intake response to optogenetic POA^Hrh1^ neuron activation. A,B) ΔTb and physical activity response to optogenetic stimulation of indicated duration (mean + SEM of 5 sequential stimulation cycles per mouse per condition; n = 5 – 10 mice per condition; males and females). ΔTb is the change from baseline (mean minus -10 to 0 minutes). Quantification (right side): ΔTb is mean of the last minute of laser on and first minute after laser end; physical activity is the first and second minute of laser on; bar is mean, circles are individual mice. C,D) Tb and physical activity response to 10 minutes of optogenetic stimulation with indicated frequency. Quantification (right side): ΔTb is Tb at 10 minutes minus Tb at 0 minutes. Physical activity is the mean during stimulation. Bar is mean, circles are individual observations. E,F) ΔTb and physical activity responses to a single 40-minute optogenetic stimulation (mean + SEM; average of 4 cycles per mouse; n = 5 – 6 mice per condition; males and females; light phase). ΔTb is the change from the -20 to 0 minutes baseline mean. Right: ΔTb and physical activity responses by hour; bar is mean, circles are individual mice. G, H) Data in A) and B) aligned by end of optogenetic stimulation. Right side, quantification of response at 50-60 minutes after end of stimulation, bar is mean, circles are individual mice. I) Food intake (chow) during one hour of optogenetic stimulation. J) 1 hr food intake (chow) after one hour of optogenetic stimulation (‘stim’) or controls (‘no stim’). A-D: p values are from one-way ANOVA with Tukey’s multiple comparison test comparing all means to each other (only relevant p values shown). E, F, I: unpaired t-test between GFP and ChR2. G, H: p values are from one-way ANOVA with Dunnett’s multiple comparisons test comparing to GFP.

**sFig 6.**
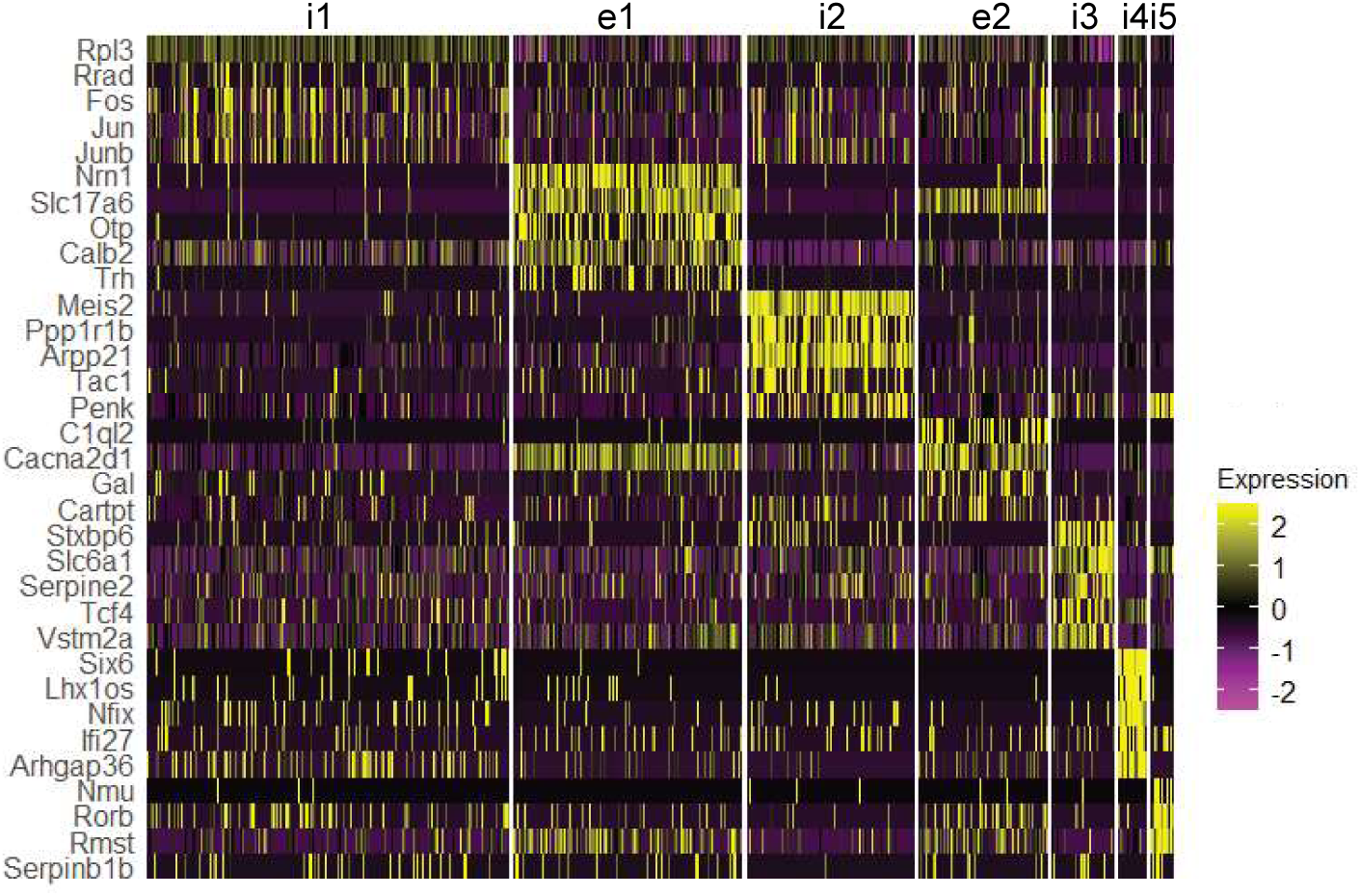
Expression profile of POA^Hrh1^ clusters. Expression profile of marker mRNAs of Hrh1-expressing clusters for the POA region. Data from ^30^

**sFig 7.**
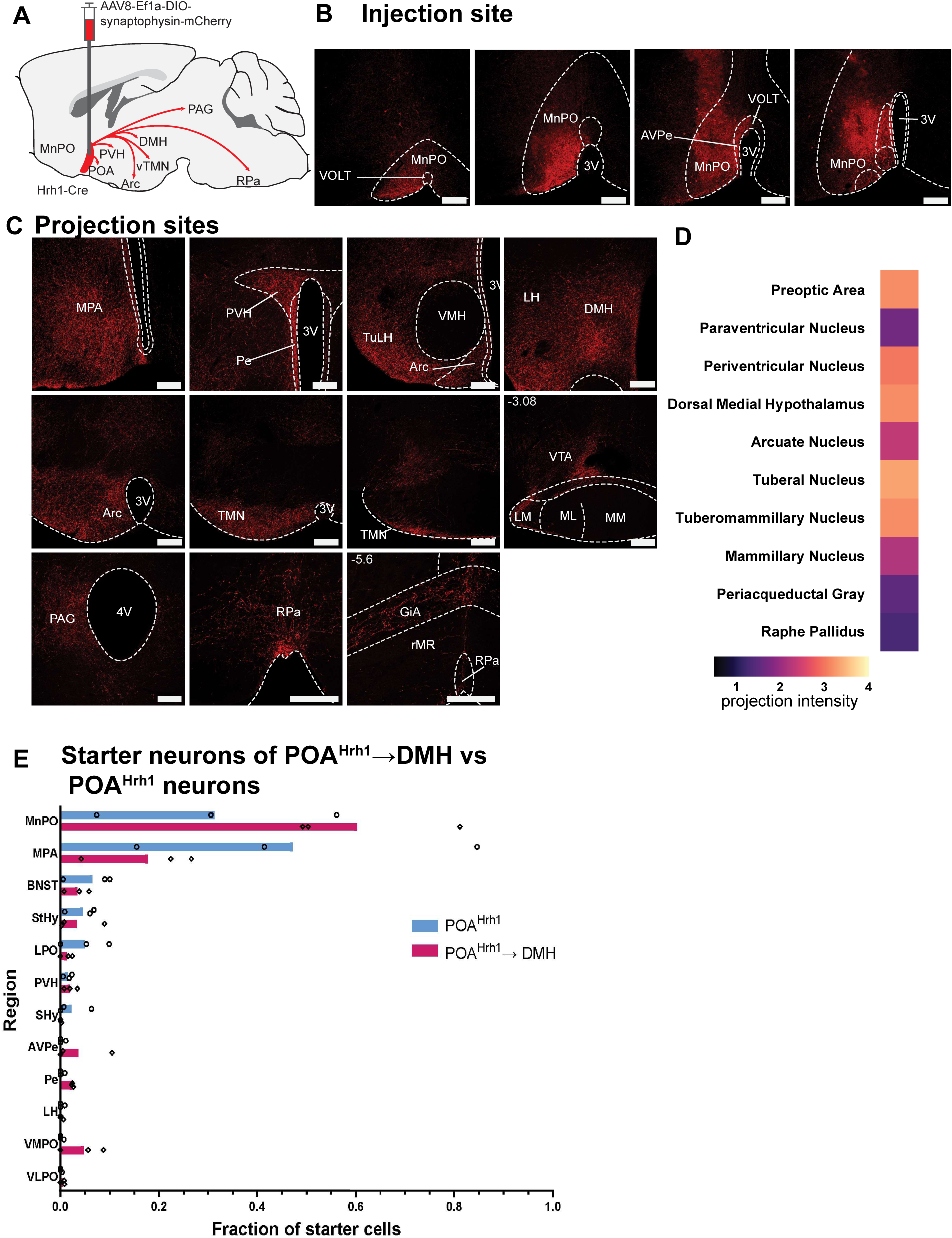
Anterograde projections of POA ^Hrh1^ neurons to wide target regions. A) Schematic of virus injection strategy. B) Example of cell body mCherry expression distribution after injection in POA. C) Examples of projection areas. Scale bar is 100 µm. D) Quantification of projection areas (n = 6 mice). E) Table with fraction of total starter neurons from Fig.4, calculated as (number of starter neurons per nucleus / total number starter neurons).

**sFig 8.**
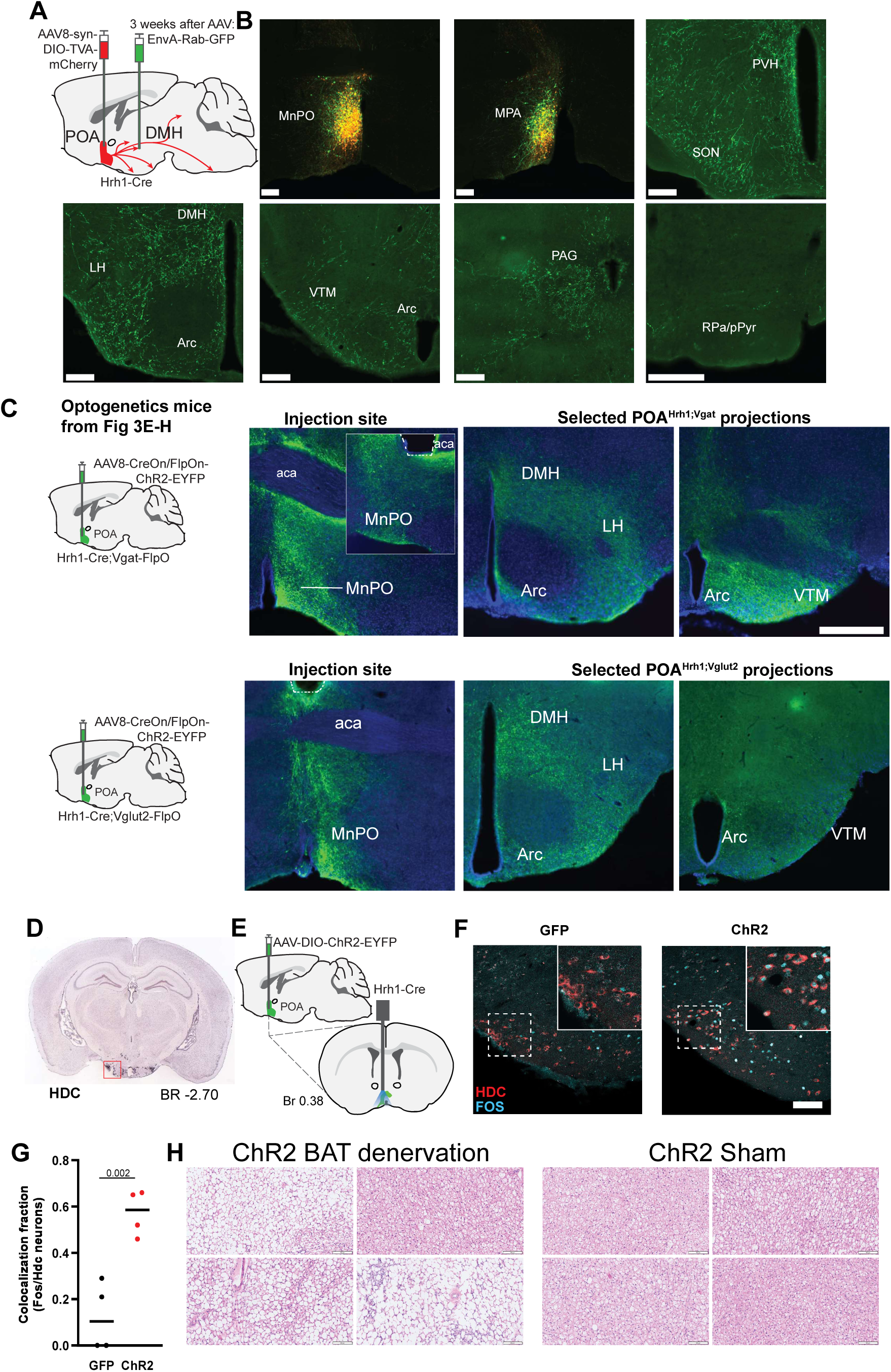
POA ^Hrh1^→DMH neurons have collaterals; POA ^Hrh1^→VTM neuron activation increases Fos in HDC neurons; BAT denervation histology. A) Schematic of virus injection strategy for tracing POA ^Hrh1^→DMH collateral projections. B) Cell bodies of POA ^Hrh1^→DMH neurons (GFP + mCherry) have collaterals to multiple hypothalamic and brainstem regions. C) Schematic of virus injection strategy (left). Examples of injection site expression of ChR2-EYFP and optic fiber placement. Examples of projection areas at two hypothalamic levels. Images are representative of 3 animals per group. Scale bar is 500 µm. D) In situ hybridization showing HDC expression neurons in the ventral tuberomammillary nucleus from Allen Brain Atlas: https://mouse.brain-map.org/experiment/show/71016663. E) Schematic of virus injection strategy and optic fiber placement for POA ^Hrh1^ neurons). F) Example of Fos expression in the VTM after 1 hour stimulation of mice expressing GFP (left) or ChR2-EYFP (right) in the POA. HDC (red) and Fos (blue) identified by immunohistochemistry. G) Quantification of Fos expression in VTM HDC neurons. P value is unpaired t-test. H) BAT histology.

**sFig 9.**
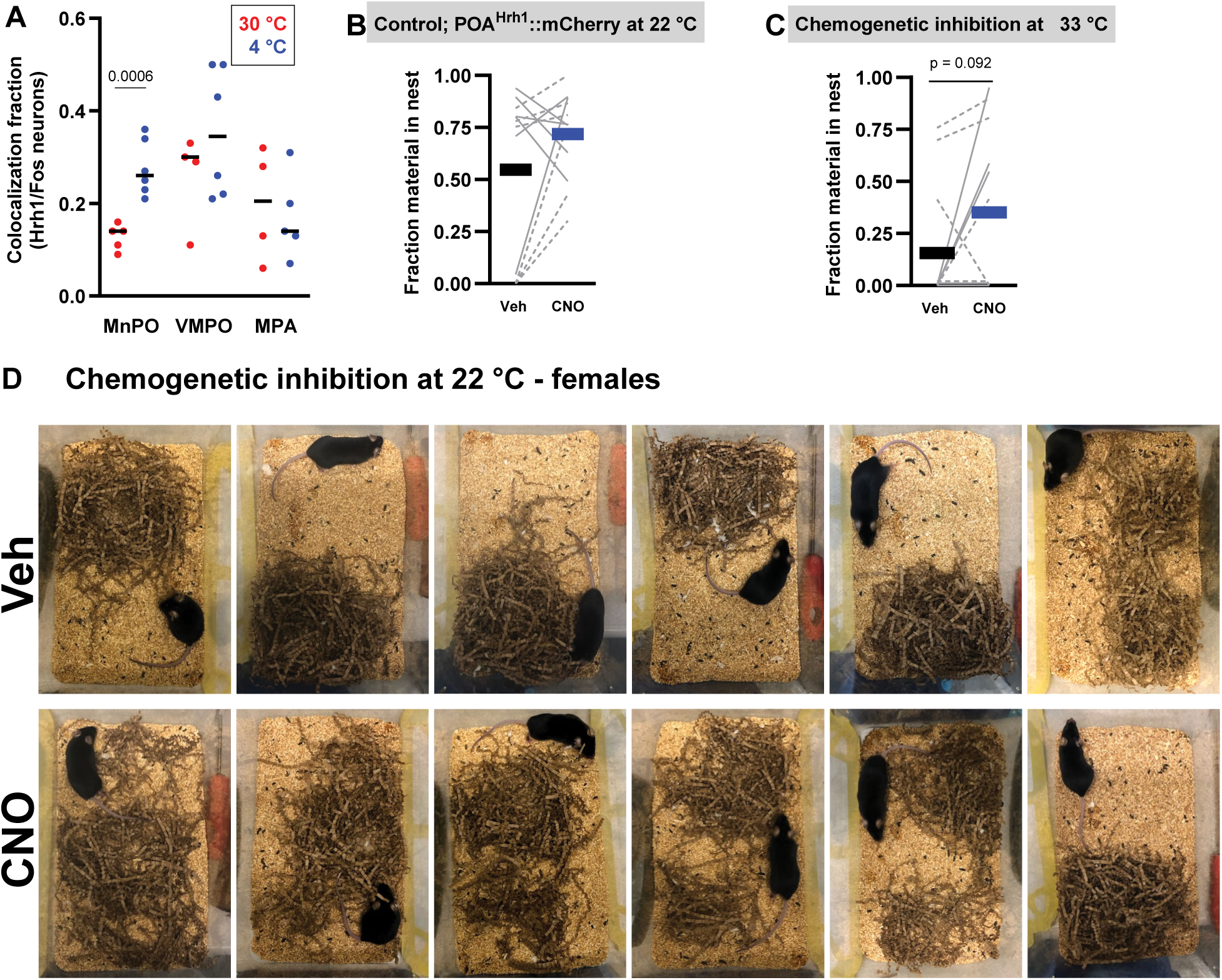
POA ^Hrh1^ neurons inhibition reduces nest-building behavior. A) Fraction of Fos neurons that expresses Hrh1 in MnPO (121 ± 19 neurons), VMPO (33 ± 7 neurons) and MPA (49 ± 10 neurons). B) Nest-building behavior in response to CNO or veh in control animals. C) Nest-building behavior in response to CNO or vehicle in at a warm ambient temperature of 33° C in POA ^Hrh1^::hM4Di mice. D) Pictures of cages at the end of the nest-building behavior assay.

## REFERENCES

1. Tan CL, Knight ZA. Regulation of Body Temperature by the Nervous System. Neuron 98, 31–48 (2018).

2. Morrison SF, Nakamura K. Central Mechanisms for Thermoregulation. Annu Rev Physiol 81, 285–308 (2019).

3. Upton BA, D’Souza SP, Lang RA. QPLOT Neurons-Converging on a Thermoregulatory Preoptic Neuronal Population. Frontiers in neuroscience 15, 665762 (2021).

4. Piñol RA, et al. Preoptic BRS3 neurons increase body temperature and heart rate via multiple pathways. Cell metabolism 33, 1389–1403 e1386 (2021).

5. Osterhout JA, et al. A preoptic neuronal population controls fever and appetite during sickness. Nature 606, 937–944 (2022).

6. da Conceicao EPS, Morrison SF, Cano G, Chiavetta P, Tupone D. Median preoptic area neurons are required for the cooling and febrile activations of brown adipose tissue thermogenesis in rat. Sci Rep 10, 18072 (2020).

7. Tan CL, et al. Warm-Sensitive Neurons that Control Body Temperature. Cell 167, 47–59 e15 (2016).

8. Panula P, Yang HY, Costa E. Histamine-containing neurons in the rat hypothalamus. Proc Natl Acad Sci U S A 81, 2572–2576 (1984).

9. Watanabe T, et al. Distribution of the histaminergic neuron system in the central nervous system of rats; a fluorescent immunohistochemical analysis with histidine decarboxylase as a marker. Brain Res 295, 13–25 (1984).

10. Wada H, Inagaki N, Itowi N, Yamatodani A. Histaminergic neuron system in the brain: distribution and possible functions. Brain Res Bull 27, 367–370 (1991).

11. Ericson H, Watanabe T, Kohler C. Morphological analysis of the tuberomammillary nucleus in the rat brain: delineation of subgroups with antibody against L-histidine decarboxylase as a marker. J Comp Neurol 263, 1–24 (1987).

12. Haas HL, Sergeeva OA, Selbach O. Histamine in the nervous system. Physiol Rev 88, 1183–1241 (2008).

13. Haas H, Panula P. The role of histamine and the tuberomamillary nucleus in the nervous system. Nat Rev Neurosci 4, 121–130 (2003).

14. Arrigoni E, Fuller PM. The Role of the Central Histaminergic System in Behavioral State Control. In: The Functional Roles of Histamine Receptors (eds Yanai K, Passani MB). Springer International Publishing (2022).

15. Green MD, Cox B, Lomax P. Histamine H1- and H2-receptors in the central thermoregulatory pathways of the rat. J Neurosci Res 1, 353–359 (1975).

16. Hong ST, et al. Histamine and its receptors modulate temperature-preference behaviors in Drosophila. J Neurosci 26, 7245–7256 (2006).

17. Lundius EG, Sanchez-Alavez M, Ghochani Y, Klaus J, Tabarean IV. Histamine influences body temperature by acting at H1 and H3 receptors on distinct populations of preoptic neurons. J Neurosci 30, 4369–4381 (2010).

18. Dong H, et al. Genetically encoded sensors for measuring histamine release both in vitro and in vivo. Neuron 111, 1564–1576 e1566 (2023).

19. Gordon CJ. Thermal physiology of laboratory mice: Defining thermoneutrality. Journal of Thermal Biology 37, 654–685 (2012).

20. Refinetti R. Circadian rhythmicity of body temperature and metabolism. Temperature (Austin) 7, 321–362 (2020).

21. Skop V, et al. Beyond day and night: The importance of ultradian rhythms in mouse physiology. Molecular metabolism 84, 101946 (2024).

22. Scammell TE, Arrigoni E, Lipton JO. Neural Circuitry of Wakefulness and Sleep. Neuron 93, 747–765 (2017).

23. Futagawa A, Tsuneoka Y, Lazarus M, Oishi Y. Comprehensive mapping of histamine H(1) receptor mRNA in the mouse brain. J Comp Neurol 532, e25622 (2024).

24. Kinnunen A, Lintunen M, Karlstedt K, Fukui H, Panula P. In situ detection of H1-receptor mRNA and absence of apoptosis in the transient histamine system of the embryonic rat brain. J Comp Neurol 394, 127–137 (1998).

25. Piñol RA, et al. Brs3 neurons in the mouse dorsomedial hypothalamus regulate body temperature, energy expenditure, and heart rate, but not food intake. Nat Neurosci 21, 1530–1540 (2018).

26. Alexander GM, et al. Remote control of neuronal activity in transgenic mice expressing evolved G protein-coupled receptors. Neuron 63, 27–39 (2009).

27. Skop V, et al. The metabolic cost of physical activity in mice using a physiology-based model of energy expenditure. Molecular metabolism 71, 101699 (2023).

28. Masaki T, et al. Involvement of hypothalamic histamine H1 receptor in the regulation of feeding rhythm and obesity. Diabetes 53, 2250–2260 (2004).

29. Michael NJ, et al. Melanocortin regulation of histaminergic neurons via perifornical lateral hypothalamic melanocortin 4 receptors. Molecular metabolism 35, 100956 (2020).

30. Moffitt JR, et al. Molecular, spatial, and functional single-cell profiling of the hypothalamic preoptic region. Science 362, (2018).

31. Machado NLS, et al. Preoptic EP3R neurons constitute a two-way switch for fever and torpor. Nature, (2025).

32. Yu S, et al. Glutamatergic Preoptic Area Neurons That Express Leptin Receptors Drive Temperature-Dependent Body Weight Homeostasis. J Neurosci 36, 5034–5046 (2016).

33. Song K, et al. The TRPM2 channel is a hypothalamic heat sensor that limits fever and can drive hypothermia. Science 353, 1393–1398 (2016).

34. Zhang Z, et al. Estrogen-sensitive medial preoptic area neurons coordinate torpor in mice. Nat Commun 11, 6378 (2020).

35. Takahashi TM, et al. A discrete neuronal circuit induces a hibernation-like state in rodents. Nature 583, 109–114 (2020).

36. Hrvatin S, et al. Neurons that regulate mouse torpor. Nature 583, 115–121 (2020).

37. Yamaguchi H, Murphy KR, Fukatsu N, Sato K, Yamanaka A, de Lecea L. Dorsomedial and preoptic hypothalamic circuits control torpor. Current biology : CB 33, 5381–5389 e5384 (2023).

38. Harding EC, et al. A Neuronal Hub Binding Sleep Initiation and Body Cooling in Response to a Warm External Stimulus. Current biology : CB 28, 2263–2273 e2264 (2018).

39. Alcantara IC, et al. A hypothalamic circuit that modulates feeding and parenting behaviours. Nature, (2025).

40. Dimicco JA, Zaretsky DV. The dorsomedial hypothalamus: a new player in thermoregulation. Am J Physiol Regul Integr Comp Physiol 292, R47–63 (2007).

41. Douglass AM, et al. Neural basis for fasting activation of the hypothalamic-pituitary-adrenal axis. Nature 620, 154–162 (2023).

42. Miklos IH, Kovacs KJ. Functional heterogeneity of the responses of histaminergic neuron subpopulations to various stress challenges. Eur J Neurosci 18, 3069–3079 (2003).

43. Cannon B, Nedergaard J. Brown adipose tissue: function and physiological significance. Physiol Rev 84, 277–359 (2004).

44. Kataoka N, Shima Y, Nakajima K, Nakamura K. A central master driver of psychosocial stress responses in the rat. Science 367, 1105–1112 (2020).

45. Yasuda T, Masaki T, Sakata T, Yoshimatsu H. Hypothalamic neuronal histamine regulates sympathetic nerve activity and expression of uncoupling protein 1 mRNA in brown adipose tissue in rats. Neuroscience 125, 535–540 (2004).

46. Shapiro JT, Michaud NM, Crowder NA. Characterizing optogenetically mediated rebound effects in anaesthetized mouse primary visual cortex. J Physiol 603, 4609–4636 (2025).

47. Machado NLS, Saper CB. Genetic identification of preoptic neurons that regulate body temperature in mice. Temperature (Austin) 9, 14–22 (2022).

48. Skop V, et al. Mouse Thermoregulation: Introducing the Concept of the Thermoneutral Point. Cell reports 31, 107501 (2020).

49. Takahashi K, Lin JS, Sakai K. Neuronal activity of histaminergic tuberomammillary neurons during wake-sleep states in the mouse. J Neurosci 26, 10292–10298 (2006).

50. Parmentier R, et al. Role of histamine H1-receptor on behavioral states and wake maintenance during deficiency of a brain activating system: A study using a knockout mouse model. Neuropharmacology 106, 20–34 (2016).

51. Zecharia AY, et al. GABAergic inhibition of histaminergic neurons regulates active waking but not the sleep-wake switch or propofol-induced loss of consciousness. J Neurosci 32, 13062–13075 (2012).

52. Venner A, et al. Reassessing the Role of Histaminergic Tuberomammillary Neurons in Arousal Control. J Neurosci 39, 8929–8939 (2019).

53. Romeijn N, et al. Sleep, vigilance, and thermosensitivity. Pflugers Arch 463, 169–176 (2012).

54. Sotelo MI, et al. Lateral hypothalamic neuronal ensembles regulate pre-sleep nest-building behavior. Current Biology 32, 806–822.e807 (2022).

55. Li XY, et al. AGRP Neurons Project to the Medial Preoptic Area and Modulate Maternal Nest-Building. J Neurosci 39, 456–471 (2019).

56. Tossell K, et al. Somatostatin neurons in prefrontal cortex initiate sleep-preparatory behavior and sleep via the preoptic and lateral hypothalamus. Nat Neurosci 26, 1805–1819 (2023).

57. Tagawa N, et al. Activation of lateral preoptic neurons is associated with nest-building in male mice. Sci Rep 14, 8346 (2024).

58. Ambler M, Hitrec T, Wilson A, Cerri M, Pickering A. Neurons in the Dorsomedial Hypothalamus Promote, Prolong, and Deepen Torpor in the Mouse. J Neurosci 42, 4267–4277 (2022).

59. Wiater MF, Li AJ, Dinh TT, Jansen HT, Ritter S. Leptin-sensitive neurons in the arcuate nucleus integrate activity and temperature circadian rhythms and anticipatory responses to food restriction. Am J Physiol Regul Integr Comp Physiol 305, R949–960 (2013).

60. Padilla SL, et al. Kisspeptin Neurons in the Arcuate Nucleus of the Hypothalamus Orchestrate Circadian Rhythms and Metabolism. Current biology : CB 29, 592–604 e594 (2019).

61. Guzman-Ruiz MA, et al. Role of the Suprachiasmatic and Arcuate Nuclei in Diurnal Temperature Regulation in the Rat. J Neurosci 35, 15419–15429 (2015).

62. Saper CB, Lu J, Chou TC, Gooley J. The hypothalamic integrator for circadian rhythms. Trends in neurosciences 28, 152–157 (2005).

63. Jain S, et al. Melanotan II causes hypothermia in mice by activation of mast cells and stimulation of histamine 1 receptors. American Journal of Physiology-Endocrinology and Metabolism 315, E357–E366 (2018).

64. Madisen L, et al. A robust and high-throughput Cre reporting and characterization system for the whole mouse brain. Nat Neurosci 13, 133–140 (2010).

65. Szymczak-Workman AL, Vignali KM, Vignali DA. Generation of 2A-linked multicistronic cassettes by recombinant PCR. Cold Spring Harb Protoc 2012, 251–254 (2012).

66. Oh SW, et al. A mesoscale connectome of the mouse brain. Nature 508, 207–214 (2014).

67. Krashes MJ, et al. Rapid, reversible activation of AgRP neurons drives feeding behavior in mice. J Clin Invest 121, 1424–1428 (2011).

68. LaFosse PK, et al. Bicistronic Expression of a High-Performance Calcium Indicator and Opsin for All-Optical Stimulation and Imaging at Cellular Resolution. eNeuro 10, (2023).

69. Deacon RM. Assessing nest building in mice. Nature protocols 1, 1117–1119 (2006).

70. Gaskill BN, Rohr SA, Pajor EA, Lucas JR, Garner JP. Some like it hot: Mouse temperature preferences in laboratory housing. Applied Animal Behaviour Science 116, 279–285 (2009).

71. Lute B, et al. Biphasic effect of melanocortin agonists on metabolic rate and body temperature. Cell metabolism 20, 333–345 (2014).

72. Gachkar S, et al. 3-Iodothyronamine Induces Tail Vasodilation Through Central Action in Male Mice. Endocrinology 158, 1977–1984 (2017).

73. Skop V, Liu N, Guo J, Gavrilova O, Reitman ML. The contribution of the mouse tail to thermoregulation is modest. American journal of physiology Endocrinology and metabolism 319, E438–E446 (2020).

74. Vianna DML, Carrive P. Stress-induced hyperthermia is not mediated by brown adipose tissue in mice. Journal of Thermal Biology 37, 125–129 (2012).

75. Vaughan CH, Zarebidaki E, Ehlen JC, Bartness TJ. Analysis and measurement of the sympathetic and sensory innervation of white and brown adipose tissue. Methods Enzymol 537, 199–225 (2014).

76. Paxinos G, Franklin KBJ. The mouse brain in stereotaxic coordinates, Compact 3rd edn. Elsevier Academic Press (2008).

77. Lutas A, et al. State-specific gating of salient cues by midbrain dopaminergic input to basal amygdala. Nat Neurosci 22, 1820–1833 (2019).

78. Zhang SX, et al. Hypothalamic dopamine neurons motivate mating through persistent cAMP signalling. Nature 597, 245–249 (2021).

79. de Araujo Salgado I, et al. Toggling between food-seeking and self-preservation behaviors via hypothalamic response networks. Neuron 111, 2899–2917 e2896 (2023).

80. Enriquez-Traba J, et al. Dissociable control of motivation and reinforcement by distinct ventral striatal dopamine receptors. Nat Neurosci 28, 105–121 (2025).

